# Phosphorylation of the synaptonemal complex protein SYP-1 promotes meiotic chromosome segregation

**DOI:** 10.1101/211672

**Authors:** Aya Sato-Carlton, Chihiro Nakamura-Tabuchi, Stephane Kazuki Chartrand, Tomoki Uchino, Peter Mark Carlton

## Abstract

Chromosomes that have undergone crossing-over in meiotic prophase must maintain sister chromatid cohesion somewhere along their length between the first and second meiotic divisions. While many eukaryotes use the centromere as a site to maintain cohesion, the holocentric organism *C. elegans* instead creates two chromosome domains of unequal length termed the short arm and long arm, which become the first and second site of cohesion loss at meiosis I and II. The mechanisms that confer distinct functions to the short and long arm domains remain poorly understood. Here, we show that phosphorylation of the synaptonemal complex protein SYP-1 is required to create these domains. Once crossovers are made, phosphorylated SYP-1 and PLK-2 become cooperatively confined to short arms and guide phosphorylated histone H3 and the chromosomal passenger complex to the site of meiosis I cohesion loss. Our results show that PLK-2 and phosphorylated SYP-1 ensure creation of the short arm subdomain, promoting disjunction of chromosomes in meiosis I.

## Introduction

The partitioning of single haploid genomes from a replicated diploid genome in meiosis requires that the four linked chromatids of each homologous chromosome pair come apart from each other in two successive divisions. Throughout these two divisions, chromosomes are mainly held together by sister chromatid cohesion. Cohesion must therefore be released in two discrete steps, so that chromosomes remain linked between the first and second division. Organisms with monocentric chromosomes first release cohesion from chromosome arms in meiosis I, but protect cohesion at the centromere using the protein Shugoshin (Kitajima et al., 2004); cohesion at the centromere is only released in the second division. Many organisms including *Caenorhabditis elegans (C. elegans)* have holocentric chromosomes (reviewed in Dernburg, 2001; and Melters et al., 2012), in which kinetochores are not restricted to a single locus but instead spread over the entire length of the chromosome. This arrangement presents a challenge to the two-step loss of chromosome cohesion, since predefined centromere and chromosome arm domains do not exist. In *C. elegans* meiosis, two-step cohesion loss is achieved by the facultative creation on each chromosome of two functionally distinct domains separated by the single crossover (CO) (Martinez-Perez et al., 2008). Since the crossover has a reliably off-center position (Barnes et al., 1995), these domains have different lengths, and are termed the short arm, which loses cohesion in meiosis I (MI), and the long arm, which retains cohesion until meiosis II (MII) (Lui and Colaiácovo, 2013). The mechanisms that determine the functional state of these domains in a length-dependent manner are not understood.

The synaptonemal complex (SC) is a macroassembly that plays critical roles in holding homologous chromosomes together (reviewed in Zickler and Kleckner, 1999) and ensuring the correct distribution of crossovers (Hayashi et al., 2010) during meiotic prophase. The SC consists of axial elements, located on the long axis of each replicated chromosome, and the central element which bridges the two axial elements (called lateral elements after synapsis). Previous studies have shown that several SC proteins disassemble asymmetrically from either short or long arms in diplotene and diakinesis, the prophase substages immediately before the first meiotic division. In wild-type animals, all the central element proteins (SYP-1, -2, -3 and -4) disassemble from the long arms of bivalents in diplotene, remain on short arms through early diakinesis, and disappear completely by the -2 position (proceeding distally from the spermatheca, oocyte precursors in diakinesis are designated as stage -1, -2, -3, etc. oocytes) (Nabeshima et al., 2005). Conversely, two of the four axial element proteins (HTP-1 and HTP-2) disassemble from short arms in diplotene and remain on long arms in diakinesis, while the remaining two (HTP-3 and HIM-3) persist on both short and long arms (Martinez-Perez et al., 2008). This asymmetric disassembly depends on the activity of the Polo-like kinase PLK-2 (Harper et al., 2011). PLK-2 first localizes to the meiotic pairing centers (PCs), DNA sequences that promote pairing in *cis* (Herman and Kari, 1989; McKim et al., 1993; Villeneuve, 1994; MacQueen et al., 2002; Phillips et al., 2005) in the leptotene/zygotene transition zone (TZ), then relocalizes to the SC from early pachytene. PLK-2 itself becomes enriched on the short arm at late pachytene (Pattabiraman et al., 2017). Although PLK-2 localization to the SC was shown to be dependent on SYP-1 (Harper et al., 2011), PLK-2 becomes confined to short arms earlier than SYP-1 does (Pattabiraman et al., 2017), leaving the mechanism of PLK-2 recruitment to the SC unexplained.

Previous studies have found SC-interacting proteins that also localize to chromosomes asymmetrically as SC proteins disassemble from either short or long arms. The protein LAB-1 forms a complex with lateral element proteins and protects cohesion on long arms at meiosis I (de Carvalho et al., 2008). LAB-1 localizes to the entire length of the SC from the TZ through pachytene and becomes confined to long arms in diplotene as SC components disassemble (de Carvalho et al., 2008). LAB-1 binds to PP1 (Protein Phosphatase 1), represented by orthologs *gsp-1* and *gsp-2* in *C. elegans,* and promotes the activity of GSP-1/2 (PP1) on the SC, antagonizing Aurora B kinase (de Carvalho et al., 2008; Tzur et al., 2012). Aurora B (*C. elegans* AIR-2) functions in a protein complex called the CPC (chromosomal passenger complex) together with INCENP (ICP-1), Borealin (CSC-1) and Survivin (BIR-1), which together play a crucial role in triggering cohesin cleavage during mitosis and meiosis (Carmena et al., 2012, 2014). The CPC also plays multiple roles in activating the spindle assembly checkpoint and destabilizing erroneous microtubule attachment to the kinetochore to ensure correct orientation of chromatids at cell division. Previous studies in *C. elegans* have shown that AIR-2 (Aurora B) localizes to short arms right before the meiosis I division, and to the interface between sister chromatids before the meiosis II division (Rogers et al., 2002; Kaitna et al., 2002). AIR-2 phosphorylates the meiotic cohesin REC-8 to trigger cohesin removal and recruits spindle assembly checkpoint proteins (Kaitna et al., 2002; Rogers et al., 2002; Dumont et al., 2010). Since the CPC dictates the site of cohesion loss and chromosome separation, its localization is strictly regulated by multiple feedback loops (Carmena et al., 2012). In diverse eukaryotic cells, two histone marks, phosphorylated histone H3 threonine 3 (H3T3ph) and phosphorylated H2A threonine 120 (H2AT120ph) are bound by the CPC and CPC-interacting Shugoshin in mitosis (Kelly et al., 2010; Yamagishi et al., 2010). H3T3 is phosphorylated by Haspin kinase (Wang et al., 2010, 2011), and this is counterbalanced by the phosphatase activity of PP1 (Qian et al., 2011) while H2AT120 is phosphorylated by Bub1 kinase (Kawashima et al., 2010). Thus, chromatin carrying both phosphorylated histones functions as a docking site for the CPC. While the mechanisms localizing the CPC to centromeres during mitosis in monocentric organisms have been well-studied, how the CPC localizes to meiotic chromosomes in holocentric organisms is not well understood.

Post-translational modification of the SC has been reported as a major aspect of its regulation (Jordan et al., 2012; Fukuda et al., 2012; Leung et al., 2015; Gao et al., 2016; Nadarajan et al., 2017). Here we report a role for C-terminal phosphorylation of SYP-1 in establishing the functions of the short and long arms in stepwise cohesion loss in meiosis. Phosphorylated SYP-1 promotes PLK-2 localization to the SC, facilitating its departure from the PCs and progression of meiotic prophase. Upon crossover formation, PLK-2 is required to enrich phosphorylated SYP-1 at the short arm, which in turn leads to restriction of PLK-2 itself to the short arm as well. Phosphorylation of SYP-1 precedes and is required for the asymmetric disassembly of SC components in late prophase, and for the enrichment of CPC-recruiting histone marks on the short arms. Loss of SYP-1 phosphorylation therefore prevents the formation of asymmetric chromosome domains, leading ultimately to the mislocalization of the CPC and failures of the first meiotic division. This work establishes SYP-1 phosphorylation as a key upstream factor in the specification of chromosomal domains important for meiotic chromosome segregation.

## Results

### C-terminal phosphorylation of SYP-1 is required for meiotic competence

To gain insight into the possible roles of SC phosphorylation, we performed a phosphoproteomics analysis using mass spectrometry of phosphoprotein-enriched protein lysates from adult *C. elegans*, in which roughly half of the cells are oocyte precursor cells in meiotic prophase. We identified 12 phosphorylation sites at the C-terminus of SYP-1 (Figure 1A, **Supplemental Table S1**), 10 of which show conservation in other *Caenorhabditis* species (Figure S1A). To test whether phosphorylation of these residues is important for SYP-1 function in meiosis, we constructed a strain expressing a transgene with a non-phosphorylatable SYP-1 allele termed 12A, with all 12 potential phosphoresidues changed to alanine. In a background lacking the endogenous *syp-1* gene, the 12A allele displayed reduced viability (59.5% viable), and a *H*igh *i*ncidence of *m*ale progeny (Him) phenotype (6%) (Figure 1B). The Him phenotype reflects meiotic X chromosome nondisjunction in XX hermaphrodite self-progeny, since worms with a single X (XO) develop as males. The *syp-1(me17)* null allele shows severely reduced viability (5% viable) and 38% males in the surviving self-progeny (MacQueen et al., 2002), indicating that *syp-1(12A)* is a partial loss of function allele. A wild-type SYP-1 transgene integrated at the same site as the 12A allele restored fertility to wild-type levels in the presence of *syp-1*(*me17*), showing that the defects in the 12A allele are specific to the introduced mutations. In oocytes undergoing the first meiotic division, we found anaphase chromosome bridges between separating chromatin masses (Figure S1B) in 12A and T452A mutants, but not in wild-type animals: 36% of anaphase I nuclei had chromosome bridges (n=11 for both 12A and T452A) whereas no bridges were detected in the wild type (n=15). This observation suggests that phosphorylation of the SYP-1 C-terminus is required for proper chromosome segregation during meiosis.

**Figure 1:**
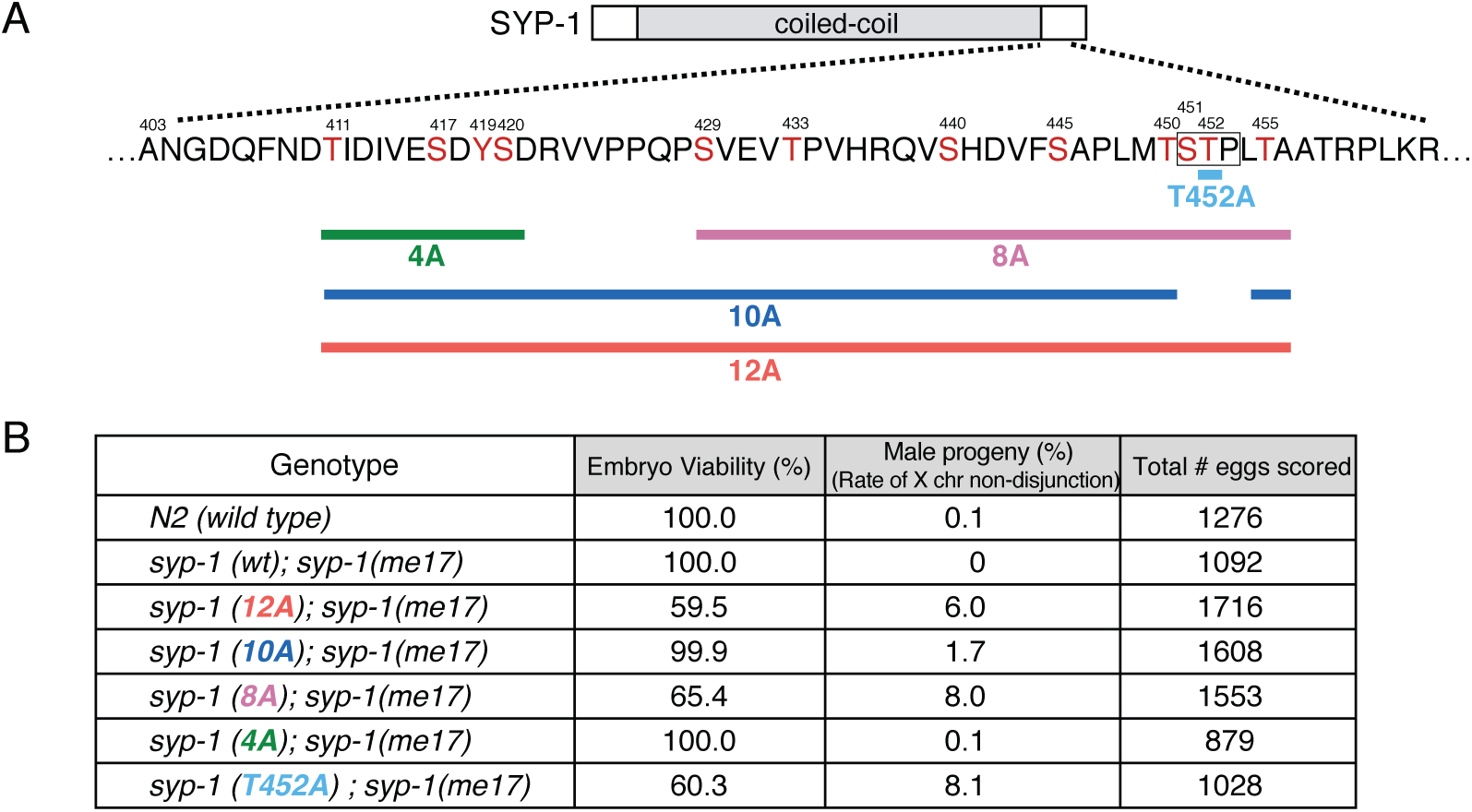
Phosphorylation of the C-terminus of SYP-1 is required for correct chromosome segregation in meiosis. (A) SYP-1 phosphorylation sites identified by mass spectrometry, and schematic diagram of a series of *syp-1* phospho-mutants. Phosphorylation sites are depicted in red, and the conserved PBD-binding motif [STP] is boxed. For each mutant, the indicated Ser/Thr/Tyr residues are converted to Ala. (B) Percentage of viability and males among the self-progeny of worms with the indicated genotypes.

### Phosphorylation of SYP-1 Threonine 452 is crucial to the function of SYP-1

To dissect the role of the SYP-1 phosphorylation sites, we made a series of transgenic worms expressing SYP-1 proteins with non-phosphorylatable mutations. First, we made *syp-1(4A)* mutants in which the first 4 phosphosites were converted to alanine, as well as *syp-1(8A)* mutants in which the last 8 phosphosites were converted to alanine. The transgenic line expressing SYP-1(4A) showed no progeny inviability or Him phenotype, whereas the line expressing SYP-1(8A) showed inviability and male production comparable to the line expressing SYP-1(12A) (Figure 1B). The most strongly conserved region within these 8 residues contains a putative S-[pS/pT]-P/X Polo-box domain (PBD)-binding motif, in which phosphorylation of the central Ser/Thr residue by a priming kinase (Elia et al., 2003) permits subsequent binding of Polo-like kinase. Our phosphoproteomics analysis detected phosphorylated Thr^452^ in this putative PBD-binding motif (Ser^451^-Thr^452^-Pro^453^) at a higher level (55 peptide counts total) than the other phosphosites (ranging from 1 to 24 counts) (**Supplemental Table S1**). To test whether this putative PBD-binding motif is important for SYP-1 function, we generated *syp-1(T452A)* mutants in which Thr^452^ in the PBD-binding motif was converted to alanine, as well as a *syp-1(10A)* mutant in which the 10 phosphosites outside the PBD-binding motif were converted to alanine. The *syp-1(T452A)* mutants showed progeny inviability and male production similar to the 12A and 8A mutants. In contrast, *syp-1(10A)* mutants were fully viable, and displayed only a weak Him phenotype (Figure 1B). This suggests that phosphorylation at the PBD-binding motif is crucial for SYP-1 function.

### Phosphorylated SYP-1 concentrates on short arms of chromosomes after crossovers are formed

To investigate the timing and location of SYP-1 phosphorylation, we generated two different phospho-specific antibodies: one against a peptide from SYP-1’s C-terminus with phosphothreonine at Thr452 (1-phos antibody), and another against the same peptide phosphorylated at three residues Thr450, Thr452 and Thr455 (3-phos antibody). Immunofluorescence staining showed similar patterns with both antibodies, but higher background with the 3-phos antibody; thereafter, 1-phos antibody was consistently used to detect phosphorylated SYP-1. The specificity of the antibody to phospho-SYP-1 was confirmed by lack of staining in T452A mutants (Figure S1C). Using this antibody, we first observed phosphorylated SYP-1 signals in the TZ, coextant with pan-SYP-1 signals along the entire length of the SC (Figure 2A). As meiocytes progress into late pachytene, and the crossover designation marker COSA-1 starts to appear, SYP-1-phos staining concentrates on short arms and decreases on long arms, while pan-SYP-1 signals are still detected along the entire SC (Figure 2B,C). In diplotene, SYP-1-phos signal intensity increases further on short arms, while pan-SYP-1 starts to disassemble from long arms, as previously shown (Martinez-Perez et al., 2008). The observation that SYP-1-phos signals begin to accumulate on short arms shortly after GFP::COSA-1 foci are detected suggested that crossovers trigger the confinement of SYP-1-phos to short arms. To test this hypothesis, we examined SYP-1-phos in gonads of *spo-11(me44)* mutants, which cannot initiate programmed double-strand breaks (DSBs) and therefore lack COs (Dernburg et al., 1998). In the absence of SPO-11, SYP-1-phos signals start to appear in the TZ but remain along the entire length of the SC through late pachytene and eventually disappear from the SC at the end of pachytene or diplotene (Figure 2D), providing evidence that double-strand breaks are required for the asymmetric distribution of phosphorylated SYP-1.

**Figure 2:**
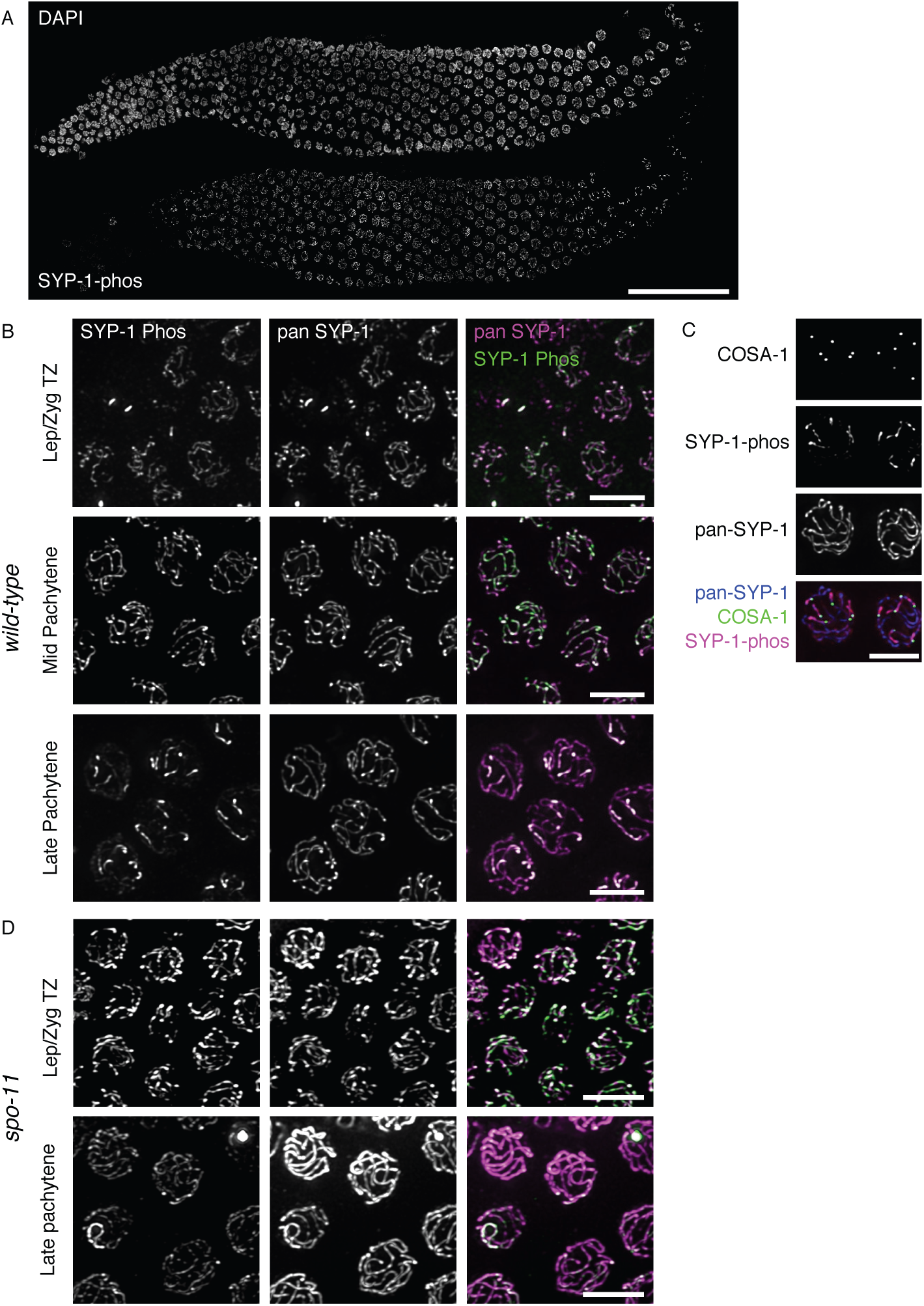
Phosphorylated SYP-1 localizes to the entire SC in early prophase, then becomes progressively restricted to the short arm. (A) A wild-type gonad stained with DAPI (top) and antibodies against phosphorylated SYP-1 (bottom). Scale bar, 50µm. (B) High-magnification images of SYP-1-phos (green) and pan-SYP-1 (magenta) staining from the indicated gonad regions showing SYP-1-phos on chromosomal subdomains in late pachytene. (C) Short-arm restriction of SYP-1-phos (magenta) in late pachytene nuclei shown by GFP::COSA-1(green) marking of crossover designation sites; pan-SYP-1 is shown in blue. Scale bar, 5µm. (D) SYP-1-phos and pan-SYP-1 staining in *spo-11 (me44)* mutant gonads. Scale bar, 5µm.

The confinement of SYP-1-phos signals to short arms was reminiscent of the previously-reported localization of PLK-2. PLK-2 first localizes to the PCs during zygotene by binding to HIM-8 and ZIM proteins, then relocates to the entire length of the SC in pachytene in a SYP-1-dependent manner, and later becomes confined to the short arms in a CO-dependent manner (Harper et al., 2011; Labella et al., 2011; Pattabiraman et al., 2017). We examined *syp-1(12A)* and *syp-1(T452A)* mutants and found that PLK-2 remained at PCs longer, eventually disappearing from the PCs in the region corresponding to late pachytene but failing to relocate to the SC (Figure 3). This result strongly suggests that the phosphorylated PBD-binding motif of SYP-1 is required for PLK-2 localization to the SC in pachytene and subsequent stages. In contrast, in *syp-1(10A)* mutants, PLK-2 also remained at PCs longer but eventually did colocalize weakly to the SC in late pachytene (Figure S1D,E). Since PBD-binding motifs must be phosphorylated at the central Thr^452^ to enable PLK-2 recruitment, the conversion of the nearby Thr^450^ to alanine in the 10A mutant might lower priming phosphorylation at Thr^452^ or confer weaker affinity to PLK-2 itself.

**Figure 3:**
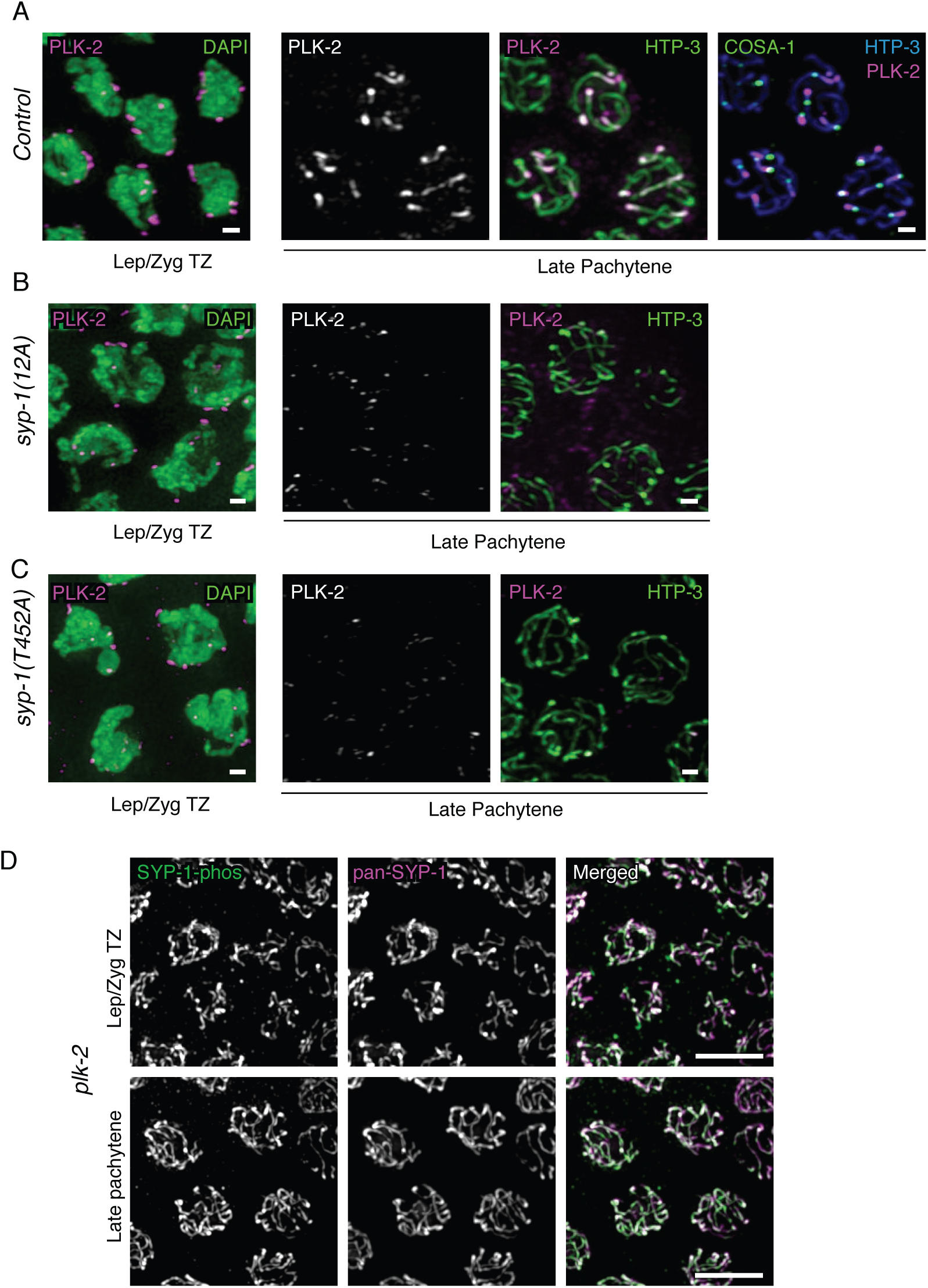
Phosphorylation of SYP-1 at Thr452 (PBD-binding motif) is required for PLK-2 relocalization from the PC to the SC. (A) *left:* Wild-type (N2) oocyte precursor cells in the leptotene/zygotene TZ immunostained with DAPI (green) and PLK-2 (magenta); *right:* oocyte precursor cells in late pachytene in control *gfp::cosa-1* animals immunostained with PLK-2 (magenta) and HTP-3 (green), or PLK-2 (magenta), HTP-3 (cyan) and COSA-1::GFP (green) in the rightmost column. (B), as (A) but for *syp-1(12A); syp-1(me17)* mutants, and without COSA-1 staining. (C), as (B) but for *syp-1(T452A) syp-1(me17)* mutants. Immunostaining shows PLK-2 relocalization from PCs to the SC (short arms) in control animals but not in the *syp-1* mutants. Scale bars, 1µm. D) SYP-1-phos and pan-SYP-1 localization in *plk-2(ok1936)* mutant gonads. Scale bars, 5µm. In all images, color of text label indicates color of the corresponding signal in merged images.

To examine whether PLK-2 acts strictly downstream of SYP-1, we visualized SYP-1-phos in *plk-2* mutants. The *plk-2(ok1936)* null mutant shows delayed synapsis, but more than 60% of chromosomes do achieve homologous pairing and synapsis, and many crossovers are made by late pachytene (about 19% of meiocytes achieve 6 COs per nucleus based on the quantification of DAPI bodies) (Harper et al., 2011; Labella et al., 2011) leading to 33.8% viable progeny (8.4% male) (Harper et al., 2011). The presence of meiocytes carrying COs in *plk-2 (ok1936)* mutants let us examine the confinement of phosphorylated SYP-1 to short arms in this mutant. We found that SYP-1 phosphorylation starting from the TZ is robustly detected in *plk-2(ok1936)* mutants, but persists over the entire length of the SC in late pachytene, never becoming confined to short arms (Figure 3D). This observation suggests the existence of a feedback loop wherein PLK-2 is required to confine SYP-1-phos to short arms in response to CO formation, and SYP-1-phos in turn restricts PLK-2 itself to short arms.

To gain insight into the chromosome segregation defects in 12A mutants, we next examined earlier steps in meiosis: chromosome pairing, synapsis, recombination, and crossover formation. Homologous chromosome pairing was assessed by staining of ZIM proteins (Phillips and Dernburg, 2006) binding the PC of chromosomes I and IV (ZIM-3), or V (ZIM-2). When chromosomes are fully homologously paired, the protein ZIM-3 which binds the PCs of chromosomes I and IV is seen as two foci, while ZIM-2, binding to chromosome V, shows a single focus per nucleus. Chromosome pairing at these sites was found to be normal in *syp-1*(12A) mutants (Figure S2A and data not shown). Next, we assessed chromosome synapsis by immunofluorescence against the SC proteins. Immunostaining showed that SYP-1(12A) protein colocalized with the axial element protein HTP-3, indicating that the non-phosphorylatable protein is expressed and correctly localized similarly to wild-type SC (Figure 4A,B). However, in *syp-1*(12A) and *syp-1(T452A)* mutants, we often observed unsynapsed chromosomes, which are positive for HTP-3 staining but missing SYP-1, in nuclei from the region corresponding to wild-type mid and late pachytene (Figure 4B,C). In contrast to *12A* and *T452A* mutants, we detected neither delayed nor partial synapsis in *10A* mutants (Figure 4D). This suggests that SYP-1 phosphorylation at the PBD-binding motif is necessary to promote timely and complete SC formation.

**Figure 4:**
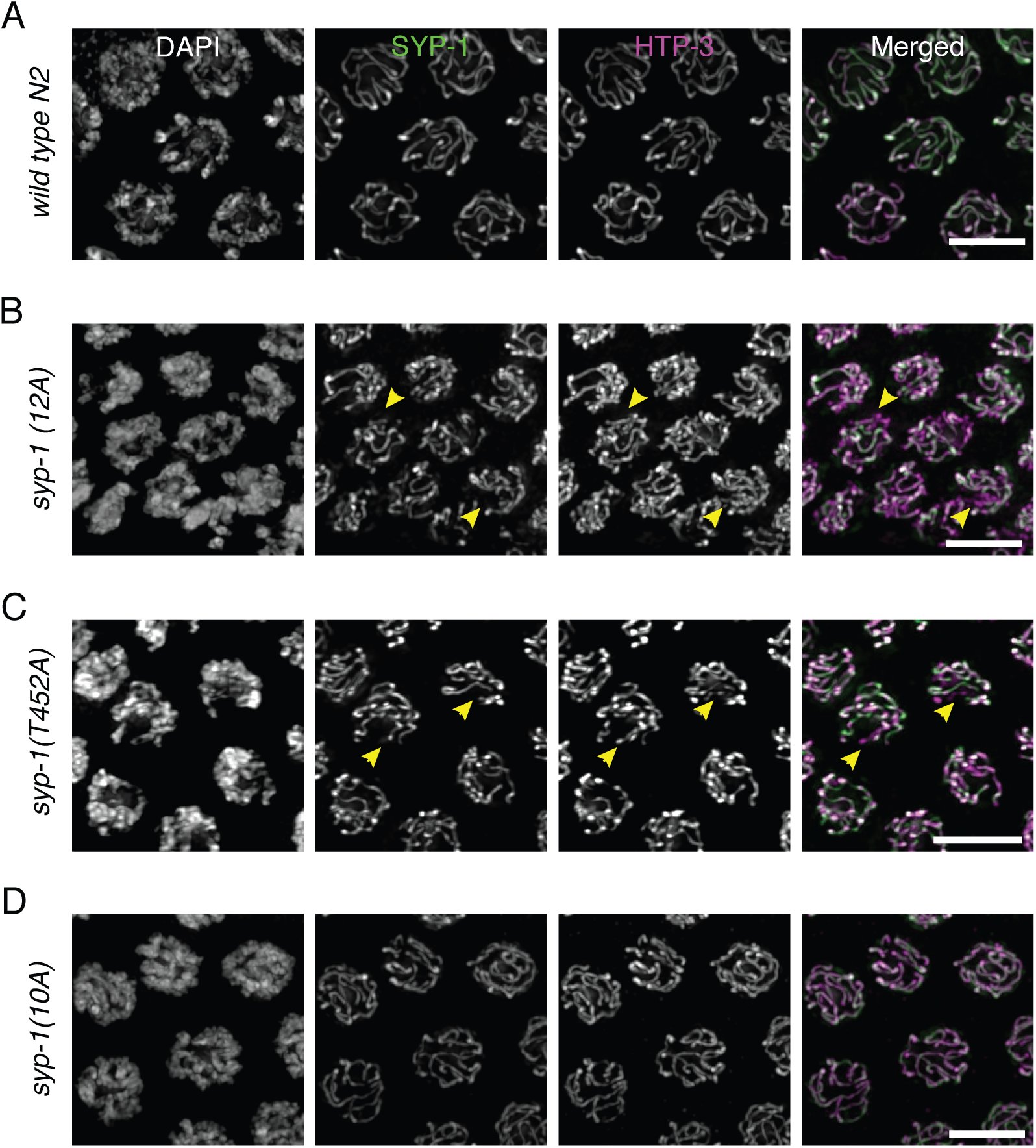
Phosphorylation of SYP-1 is required for timely completion of synapsis. Oocyte precursor cells in mid-pachytene immunostained with SYP-1 and HTP-3 antibodies in (A) wild-type, (B) *syp-1(12A); syp-1(me17),* (C) *syp-1(T452A); syp-1(me17)* and (D) *syp-1(10A); syp-1(me17)* animals. DNA is counterstained with DAPI. Unsynapsed chromosomes in 12A and T452A mutants are indicated by arrowheads. Scale bars, 5µm.

### SYP-1 phosphorylation is required for correct timing of early meiotic prophase events

We next examined the timing of progression through meiotic prophase in *syp-1* phosphorylation mutants by measuring the proportion of the gonad occupied by each substage. Meiotic nuclei move unidirectionally through the *C. elegans* gonad in a manner that allows the physical span of a substage to serve as a proxy for its duration (Hirsh et al., 1976; Jaramillo-Lambert et al., 2007). We found that gonads of both *syp-1(12A)* and *syp-1(10A)* mutants showed delayed exit from the leptotene/zygotene TZ and early pachytene stages compared to control animals (Figure 5A, B, Figure S2B, C). Previous studies have shown that meiotic checkpoints monitor the formation of CO intermediates and the status of synapsis (Bhalla and Dernburg, 2005; Carlton et al., 2006; Saito et al., 2012; Stamper et al., 2013; Rosu et al., 2013; Kim et al., 2015), extending the time spent in the TZ and early pachytene for nuclei lacking CO intermediates.

**Figure 5:**
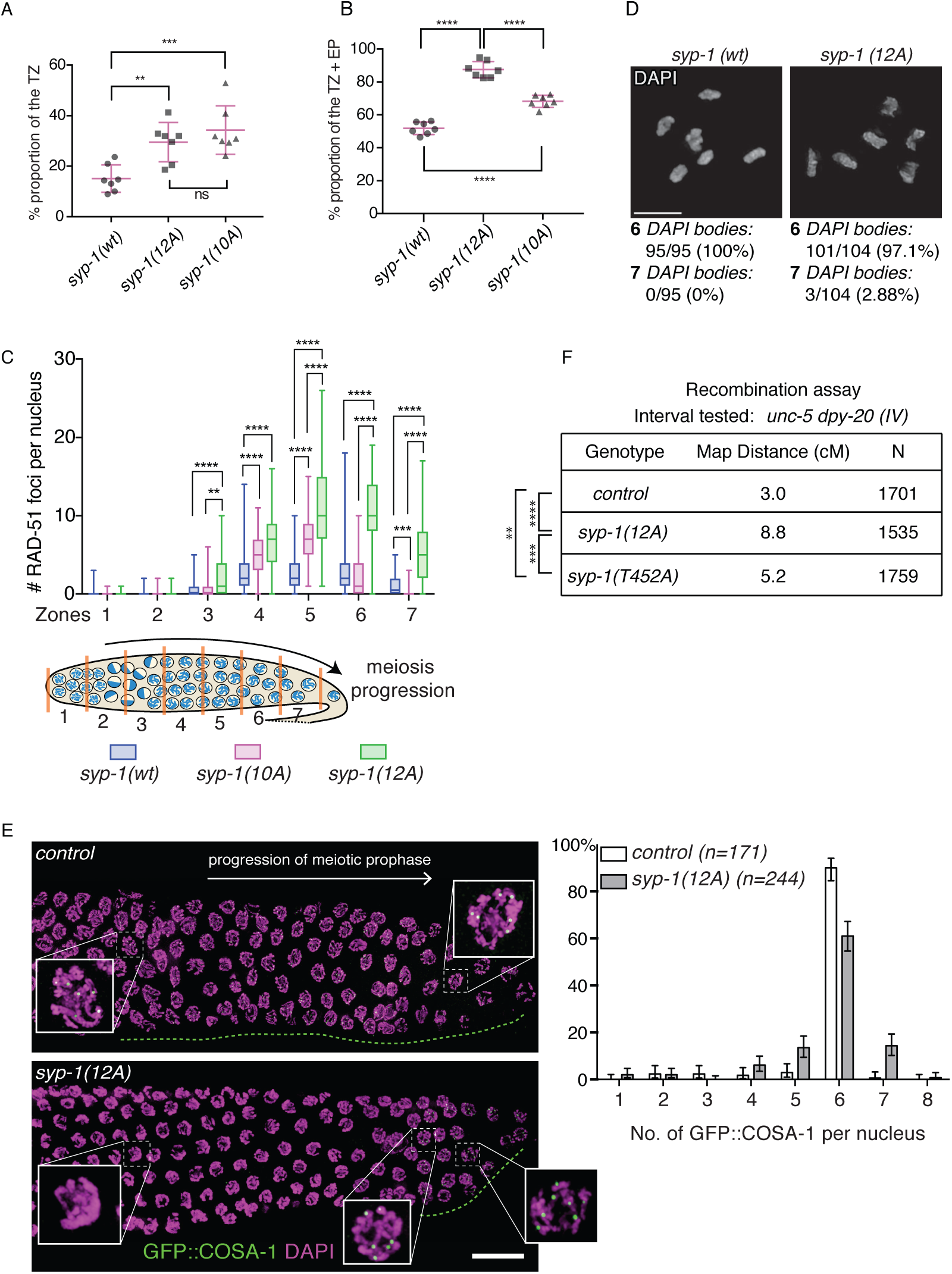
Phosphorylation of SYP-1 is required for timely progression through meiotic prophase, and affects CO distribution. (A) The proportion of TZ nuclei in *syp-1(wt); syp-1(me17)*, *syp-1(12A); syp-1(me17)* and *syp-1(10A); syp-1(me17)* animals (see also Methods and Figure S2). Seven gonads were scored for each genotype. Error bars show standard deviation. Statistical analysis was performed with a Mann-Whitney test: **p<0.01, ***p<0.001. (B) The proportion of TZ and early pachytene nuclei marked by SUN-1 Ser8-phos staining. Error bars show standard deviation. Seven gonads were scored for each genotype. ****p<0.0001, Mann-Whitney test. (C) Quantification of recombination marker RAD-51 focus counts in each of seven equal-length zones covering the TZ to late pachytene. RAD-51 focus numbers per nucleus are depicted as a box plot, with box indicating mean and quartiles, and whiskers indicating the entire range of measurements. Four gonads were scored for each genotype. The *syp-1(12A); syp-1(me17)* and *syp-1(10A); syp-1(me17)* mutants showed increased levels of RAD-51 compared to the control *syp-1(wt); syp-1(me17)*. ****p<0.0001, ***p<0.001, **p<0.01, Mann-Whitney test. (D) Representative diakinesis chromosomes from *syp-1(wt); syp-1(me17)* (left) and *syp-1(12A); syp-1(me17)* mutant (right) gonads, stained with DAPI. Statistics of DAPI body observations are shown below. A total of 95 nuclei were scored for the control and 104 nuclei scored for 12A mutants. Scale bars, 5µm. (E) *Left:* The timing of appearance of the CO designation marker GFP::COSA-1 is delayed in *syp-1(12A); syp-1(me17)* mutants. The direction of meiotic prophase progression is from left to right. The gonad region in which GFP::COSA-1 foci (green) are observed is highlighted with a green dotted line for the control *gfp::cosa-1* and *syp-1(12A) gfp::cosa-1; syp-1(me17)* gonads. DNA is counterstained by DAPI (magenta). *Right*: quantitation of COSA-1 focus numbers from three gonads for the control *gfp::cosa-1* or ten gonads for *syp-1(12A) gfp::cosa-1; syp-1(me17)* mutants. N indicates the number of total nuclei scored. Bar heights are percentages of nuclei with the given number of COSA-1 foci; error bars show 95% confidence intervals. Scale bar, 15µm. (F) Measured genetic map distances in centimorgans for the interval *unc-5---dpy-20* on chromosome IV scored on the given number (N) of *unc-5 dpy-20/+ +* (control), *syp-1(12A); unc-5 dpy-20/+ +; syp-1(me17)*, and *syp-1(T452A); unc-5 dpy-20/++; syp-1(me17)* self-progeny. ****p<0.0001, ***p=.0001484, **p=0.001336, Fisher's exact test.

When chromosomes fail to obtain CO intermediates, the cell cycle checkpoint kinase CHK-2 is activated and continues to phosphorylate PC proteins, which keeps PLK-2 bound to PCs (Kim et al., 2015). Subsequently, PC-bound PLK-2 nucleates the SUN-1/ZYG-12 nuclear envelope complex to connect chromosome ends to the cytoskeleton and promotes clustering of chromosomes, a cytological marker for the TZ and early pachytene. (Harper et al., 2011; Labella et al., 2011; Woglar et al., 2013). Gonads from 12A and 10A mutants showed an extended region of the TZ, suggesting that PLK-2 persistence at PCs maintains clustering of chromosomes in these mutants (Figure 5A). Gonads from 12A, T452A, and 10A mutants also had an extended early pachytene stage, marked by phosphorylation of the nuclear protein SUN-1 (SUN-1 Ser8-phos) (Penkner et al., 2009) (Figure 5B, S2D and data not shown for T452A). Delayed exit from early pachytene could be explained by the complete or partial inability of PLK-2 to transit from PCs to the SC. In addition, delayed synapsis leading to delayed formation of COs in 12A and T452A mutants would activate the CHK-2-mediated meiotic checkpoint (Kim et al., 2015), which promotes retention of PLK-2 by continuous phosphorylation of PC proteins, delaying the exit from early pachytene.

Next, we examined the contribution of SYP-1 phosphorylation to recombination and crossover formation, by visualizing the recombination protein RAD-51 and formation of intact bivalents. RAD-51 foci appeared with normal timing in *syp-1(12A)* and *syp-1(10A)* mutants, suggesting that the initiation of programmed DNA double strand breaks (DSBs) by SPO-11 is not perturbed in these mutants. Instead, the number of RAD-51 foci per nucleus was found to be prominently increased in mid and late pachytene (zones 4–7) in 12A, and less prominently increased in 10A mutants, compared to control animals (Figure 5C). In 12A mutants, delayed synapsis would be expected to lead to delayed homologous recombination and activate the CHK-2-mediated meiotic checkpoint, extending the window during which DSBs are generated. In contrast, 10A mutants had increased levels of DSBs without obvious delays in synapsis. This suggests that persistence of PLK-2 at PCs (or absence of PLK-2 from the SC) could suffice to maintain the DSB-generating machinery in an active state with or without activating the upstream kinase CHK-2. In addition, recent results show that PLK-1/2-mediated phosphorylation of SYP-4 is required to turn off DSB formation (Nadarajan et al., 2017). To test is SYP-1 phosphorylation is required for SYP-4 phosphorylation, we immunostained *syp-1* phosphomutants and found that SYP-4-phos signals are still detected (Figure S2F) in both 12A and T452A mutants. This suggests that the increase in RAD-51 focus numbers we observe in *syp-1(12A)* is not solely due to loss of SYP-4 phosphorylation.

To investigate whether the increase in recombination intermediates in 12A mutants could explain the observed meiotic chromosome segregation defects and 40% reduction in progeny viability as a consequence of incomplete recombination, we next examined bivalent formation in diakinesis by scoring the number of DAPI-staining bodies in late meiotic prophase. We found that 12A mutants form meiotic bivalents held together by chiasmata at nearly wild-type levels (97.1%, n=104) (Figure 5D). The fraction of diakinesis nuclei with univalents (chromosomes lacking COs) in 12A mutants was 2.9% (7 DAPI bodies: 5 bivalents and 2 univalents, n=104) while it was 0% in the wild-type (n=95). We further examined DAPI-staining bodies for the presence of intra- and inter-bivalent DNA bridges, morphological features suggestive of improperly resolved recombination intermediates (Saito et al., 2013). We found one intra-bivalent bridge out of 31 (3.2%) examined nuclei in *syp-1(12A)*, and one inter-bivalent bridge out of 11 (9.1%) examined nuclei in *syp-1(T452A)*, compared with none in 48 wild-type nuclei (0%). This suggests while *syp-1* phosphomutants show delayed meiotic progression and increased recombination, a relatively minor fraction of meiocytes suffer from improperly resolved or unrepaired recombination.

### SYP-1 phosphorylation affects crossover distribution

Previous studies have shown that SYP-1 is required for CO distribution and designation along chromosomes (Hayashi et al., 2010; Libuda et al., 2013). We tested if SYP-1 phosphorylation affects the timing and extent of CO designation by visualizing COSA-1 in late pachytene. Meiotic nuclei which obtain DSBs in mid-pachytene acquire competence to load COSA-1 at eventual CO sites (Yokoo et al., 2012). We observed a significant delay in COSA-1 appearance in 12A mutants, consistent with the observation that the TZ and early pachytene are extended and meiotic cell cycle progression is delayed. In control animals, sites marked by GFP::COSA-1 foci start to appear in late pachytene, which corresponds to about 40% of the meiotic region of the gonad (the last 40% of the proximal end). In contrast, 12A mutants showed GFP::COSA-1 foci appearing only at the very end of the gonad (last 10% of the proximal end), where meiotic nuclei finally exit early pachytene and show late pachytene chromosome morphology (Figure 5E **and** S2E). In contrast to 12A mutants, 10A mutants showed wild-type timing of GFP::COSA-1 foci appearance (Figure S2E), indicating that entrance into late pachytene is not delayed in 10A mutants. Immunofluorescence of SUN-1 Ser8-phos and GFP::COSA-1 in 10A mutant gonads suggests that although the TZ and early pachytene are prolonged in 10A mutants, entrance into late pachytene is not delayed, because of a compensatory loss of mid-pachytene (data not shown). Quantitation of GFP::COSA-1 foci revealed an increased fraction of nuclei with 7 COSA-1 foci in 12A mutants (14.3% in 12A compared to 0.6% in control), suggesting that CO designation is perturbed in the absence of SYP-1 phosphorylation. Although we detected a slightly increased number of nuclei with only 5 GFP::COSA-1 foci in 12A (13.3% in 12A compared to 2.9% in control), eventual completion of a 6th CO or apoptotic culling would result in a lower frequency of diakinesis nuclei with univalents, as we observe. To further assess the extent of crossover recombination in *syp-1* phospho-mutants, we measured the genetic distance between the *unc-5* and *dpy-20* genes on chromosome IV. In control animals, the distance was calculated as 3.0 cM, in agreement with the reference map distance of 3.44 cM (WormBase WS260). However, the 12A and T452A alleles showed significantly larger distances (Figure 5F), suggesting either an elevated crossover frequency or changes in the recombination landscape. Interestingly, the map distance measured in 12A mutants (8.8 cM) was also significantly larger than that in T452A mutants (5.2 cM), raising the possibility that the different levels of SYP-1 phosphorylation might influence CO designation capacity differently. These experiments show that loss of SYP-1 phosphorylation alters levels of synapsis and recombination as well as CO designation and distribution, which together are likely to partially contribute to the inviability observed in the *syp-1* phosphomutants.

### SYP-1 phosphorylation promotes establishment of short/long arm asymmetry

Previous studies have shown that PLK-2 plays essential roles in the establishment of short and long arm asymmetry in addition to its role in homolog pairing and synapsis (Harper et al., 2011; Labella et al., 2011; Pattabiraman et al., 2017; Nadarajan et al., 2017). The observed confinement of SYP-1-phos signals to short arms also raises the possibility that SYP-1 phosphorylation is involved in the functional distinction of the short and long arms. To test this hypothesis, we used immunofluorescence to examine protein localization on diakinesis chromosomes. Wild-type SYP-1 departs from long arms as bivalents undergo diplotene remodeling (Figure 6A), and eventually departs from short arms no later than the -3 oocyte (Figure S3). In contrast, we found that non-phosphorylatable SYP-1 as well as SYP-2 always persists on both short and long arms of diakinesis chromosomes until the -2 or -1 oocyte in the *syp-1(12A)* or *(T452A)* mutants (Figure 6B, 6C, S3D and data not shown for SYP-2). This result shows that phosphorylation of SYP-1 is required for its timely removal from long and short arms. We next visualized proteins reported to dissociate from the short arm after crossover formation: HTP-1/2, LAB-1, and COH-3/4 (Martinez-Perez et al., 2008; de Carvalho et al., 2008; Severson and Meyer, 2014). These proteins also remained on both short and long arms on at least one chromosome in the majority of -1 oocytes in *syp-1*(12A) and *syp-1*(T452A) mutants (Figure 6B, C; Figure S3A). These results indicate that C-terminal phosphorylation of SYP-1 promotes the establishment of short/long chromosome arm asymmetry, as well as the timely disassembly of SYP-1 from the SC. To further understand the function of SYP-1 phosphorylation, we created alleles with all 12 phosphosites mutated to D or E (12D or 12E), an alteration which in some cases mimics phosphorylation. However, we found that viability, male progeny production, and immunostaining phenotypes of *syp-1(12E)* and *syp-1(12D)* were indistinguishable from *syp-1(12A)*, with SYP-1 persisting on both short and long arms in diakinesis (Figures S2C, S3B, C and D). From this we conclude that SYP-1(12D) or (12E) may functionally resemble the non-phosphorylatable 12A alleles, rather than mimicking constitutively phosphorylated SYP-1.

**Figure 6:**
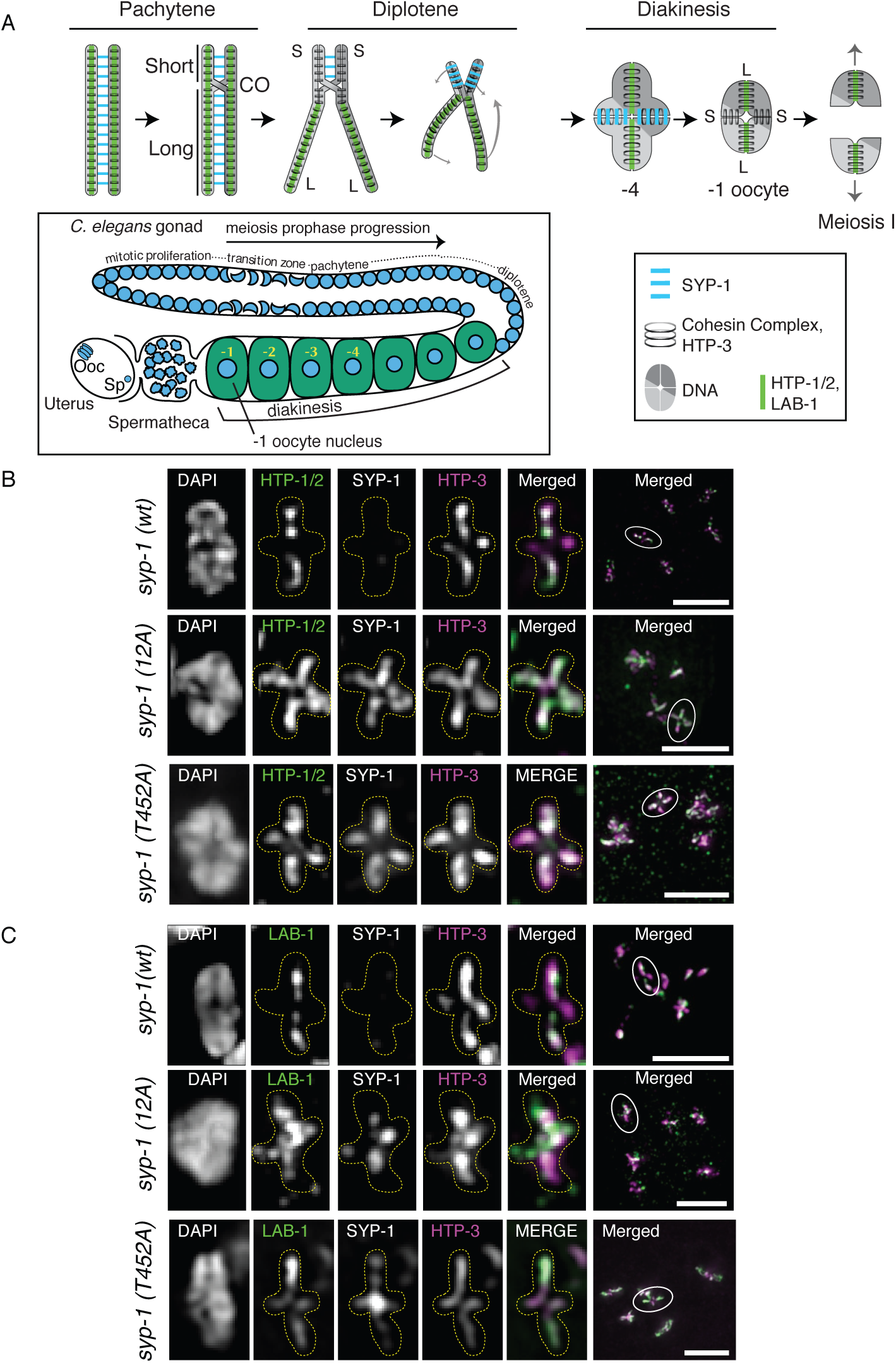
Phosphorylation of SYP-1 is required for the asymmetric localization of factors involved in chromosome segregation. (A) *Top:* Diagram of chromosome axis remodeling during meiotic prophase from pachytene through meiosis I. Cruciform axis of HTP-3/cohesin with short arm (S) and long arm (L) is shown in diakinesis. *Bottom:* Diagram of meiotic prophase substages in the *C. elegans* gonad with meiotic nuclei indicated by blue circles. Ooc: oocyte nucleus, Sp: sperm pronucleus. Diakinesis nuclei are referred to by their position relative to the spermatheca, with the nearest diakinesis nucleus being the -1 oocyte. After entering the spermatheca and becoming fertilized, the -1 oocyte nucleus goes through the meiosis I and II divisions to generate the oocyte pronucleus. (B and C) Partial Z-projection images of a representative chromosome pair in -1 or -2 oocytes with immunostaining for HTP-1/2 (green), SYP-1, and HTP-3 (magenta) (B) or LAB-1 (green), SYP-1, and HTP-3 (magenta) (C) in the *syp-1(wt); syp-1(me17)* (-1 oocyte)*, syp-1*(12A)*; syp-1(me17)* (-1 oocyte) and *syp-1(T452A); syp-1(me17)* (-2 oocyte) gonads. The 12A and T452A mutant gonad shows aberrant persistence of SYP-1 on both arms (see also Fig. S3D), and HTP-1/2 and LAB-1 on short arms, in -1 or -2 oocytes. In both B and C, representative chromosomes are presented with the color of image label indicating the color of immunostaining in the merged image. Full projection images of each nucleus are shown in the rightmost column, with the chromosome pair shown on the left encircled. The cruciform axis of HTP-3 is visible only for chromosomes lying perpendicular to the optical axis.

### The Chromosomal Passenger Complex is mislocalized in *syp-1* non-phos mutants

The arm asymmetry defects and anaphase bridges we observed in *syp-1(12A)* mutants next prompted us to examine the localization of the chromosomal passenger complex (CPC). Loss of arm asymmetry caused by *lab-1* or *htp-1* mutations has been linked to mislocalization of AIR-2 (Aurora B) and the chromosomal passenger complex (CPC), of which AIR-2 (Aurora B) is a part, leading to chromosome segregation defects (de Carvalho et al., 2008; Martinez-Perez et al., 2008). To examine whether *syp-1(12A)* mutations lead to CPC mislocalization, we looked at -1 oocytes of 12A mutants. In control animals expressing GFP::AIR-2, we observed the expected concentration of AIR-2 (Aurora B) on short arms in -1 oocytes, followed by its localization to the meiotic spindle in anaphase I, and re-localization between sister chromatids at metaphase II. In contrast, in *syp-1*(12A) mutants, AIR-2 (Aurora B) was faint and disorganized in many of the corresponding oocytes, as well as on chromosomes in metaphase/anaphase I. (Figure 7A). Similarly, the CPC component IN-CENP (ICP-1 in *C. elegans*) shows robust localization to short arms in wild-type -1 oocyte nuclei, whereas *syp-1(12*A) and *syp-1 (T452A)* mutants had significantly reduced levels of ICP-1 signals in the corresponding nuclei: 81% of nuclei showed reduced ICP-1 signals compared to the wild type while 19% of nuclei showed no ICP-1 signal on any chromosomes in 12A mutants (Figure 7B, S4). In *plk-2* mutants, an even stronger loss of ICP-1 from short arms was observed in -1 oocytes: 40% of nuclei showed reduced ICP-1 signals compared to the wild-type while 60% of nuclei showed no ICP-1 staining on any chromosomes. We noted that the magnitude of ICP-1 reduction does not correspond to the level of progeny viability in *syp-1(12A)* (60% viability) or *plk-2 (ok1936)* (34% viability) (Harper et al., 2011) mutants.

**Figure 7:**
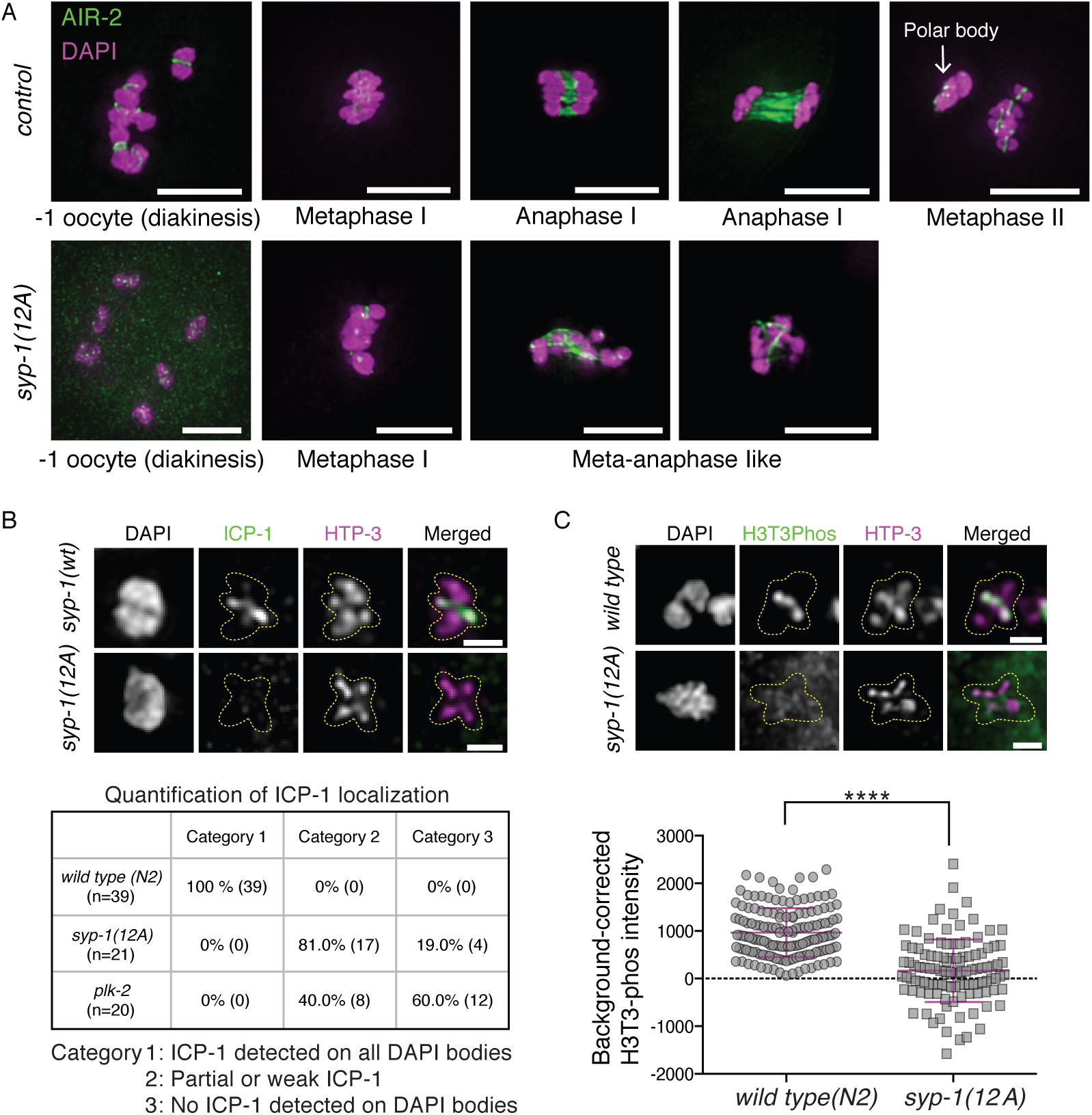
SYP-1 phosphorylation is required for correct localization of CPC components (AIR-2, ICP-1) and the CPC-guiding histone mark (H3T3ph). (A) Localization of GFP::AIR-2 (green) on meiotic chromosomes fixed and stained with DAPI (magenta) at different stages preceding and during the meiotic divisions in control (*gfp::air-2*)(top) and *gfp::air-2; syp-1(12A); syp-1(me17)* (bottom) animals. Scale bars, 5µm. (B) ICP-1 is enriched on short arms in -1 oocytes in *syp-1(wt); syp-1(me17)* animals (top) whereas some -1 oocytes lack ICP-1 localization in *syp-1(12A); syp-1(me17)* mutants (bottom) (see also whole nucleus image in Figure S4). The ICP-1 localization in -1 oocytes was categorized as follows in *syp-1(wt); syp-1(me17), syp-1(12A); syp-1(me17)* and *plk-2(ok1936)*: all 6 DAPI bodies had robust ICP-1 staining on short arms (Cat.1); some DAPI bodies had partial or weak ICP-1 staining (Cat.2); no DAPI bodies had ICP-1 staining (Cat.3). The analysis was limited to -1 oocyte nuclei carrying six bivalents. Scale bars, 1µm. (C) H3T3ph staining in -1 oocyte nuclei in N2 wild-type (top) and *syp-1(12A); syp-1(me17)* mutants (bottom); representative images are shown above quantitation (see also whole nucleus image in Figure S4). Scale bars, 1µm. In the scatter plot below, each point is a measurement of a single DAPI body. The number of DAPI bodies/points counted for H3T3ph is 151 for wild-type and 111 for *syp-1*(12A) mutants. ****p<0.0001, Mann-Whitney test.

Further examination revealed that while wild-type gonads always show ICP-1 signals on short arms from the -3 or −4 oocyte stage, some of the *syp-1(12A)* and *plk-2 (ok1936)* mutant gonads have ICP-1 signals on prometaphase I chromosomes but not diakinesis chromosomes (Figure S4A **for *plk-2***, data not shown for 12A). This suggests that ICP-1 is sometimes capable of rapidly accumulating on chromosomes upon entrance to the meiosis I division without significant prior accumulation in prophase. This is likely a result of redundant positive feed-back mechanisms that enhance CPC localization (Carmena et al., 2012), and could partially explain the significant progeny viability observed in *syp-1(12A)* and *plk-2 (ok1936)* mutants. Consistent with reduced localization of the CPC in *syp-1(12A)* mutants, levels of H3Ser10 phosphorylation, which is mediated by AIR-2 (Aurora B) kinase (Hsu et al., 2000), were also reduced in *syp-1(12A)* mutants compared to the wild type (Figure S4D). The mislocalization of the CPC could lead to failures in triggering cohesin cleavage and the spindle assembly checkpoint, and could explain the anaphase chromosome bridges and loss of viability found in *syp-1*(12A) mutants.

Since LAB-1 has been shown to promote the phosphatase activity of GSP-2 (PP1) at the SC in wild-type animals (Tzur et al., 2012), and we observed mislocalization of LAB-1 and the CPC in *syp-1(12A)* mutants, we next examined the localization of phosphorylated H3T3, which has been shown to recruit the CPC and is a substrate of PP1 in other model organisms (Qian et al., 2011). In wild-type germlines, H3T3ph signals appeared on the short arm of diakinesis chromosomes from the -3 to -4 oocyte stage. In contrast, H3T3ph was strikingly reduced or absent from short arms in -1 oocytes in *syp-1*(12A) mutants (Figure 7C **and** S4C). This suggests that in 12A mutants, LAB-1 mislocalizes to both long and short arms, promoting dephosphorylation of H3T3 via GSP-2 (PP1) on the entire chromosome.

To confirm that PP1 dephosphorylates H3T3ph in *C. elegans* meiosis, we immunostained H3T3ph in worms homozygous for the *gsp-2*(PP1) deletion allele *tm301*. In contrast to the short arm-specific H3T3ph staining in wild-type, *gsp-2*(PP1) mutants showed H3T3 phosphorylation over the entire chromosome from diakinesis onward (Figure 8A). This suggests that dephosphorylation of H3T3 by PP1 is likely to be conserved in *C. elegans*. Similarly to H3T3ph staining, ICP-1 was present all over chromosomes in *gsp-2 (PP1)* mutants by immunofluorescence (Figure 8B).

**Figure 8:**
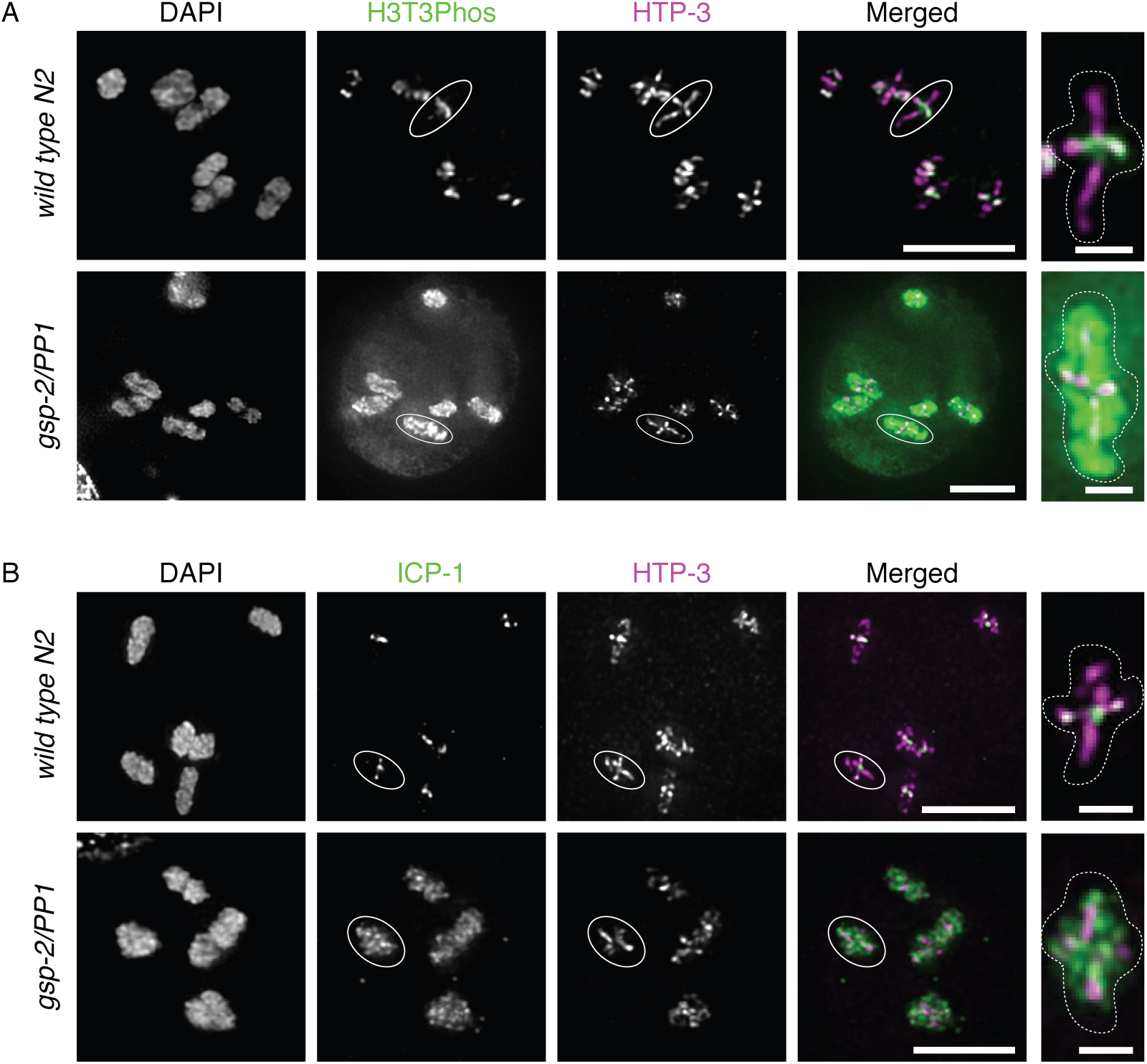
GSP-2/PP1 restricts the CPC-guiding histone mark H3T3ph and ICP-1 to short arms. H3T3ph immunostaining shown in green (A) or ICP-1 immunostaining shown in green (B) in -1 oocytes in wild-type (top) and *gsp-2(tm301)* mutants (bottom). Short and long arms are marked by HTP-3 staining (magenta). Representative chromosomes with cruciform HTP-3 staining indicated by circles are shown in the rightmost panels. Scale bars: 5µm in left panels, 1µm in the rightmost panels.

Taken together, the high incidence of defects in the asymmetric disassembly of SC components and compromised CPC localization likely explain the bulk of reduced viability in *syp-1(12A)* and *(T452A)* mutants, by causing chromosome segregation problems at meiosis I. The loss of short/long arm asymmetry in the non-phosphorylatable SYP-1 alleles and the dynamic localization of SYP-1-phos signals are two independent lines of evidence that strongly suggest that phosphorylation of SYP-1 is a key element of the molecular cascade that establishes domains of successive cohesion loss *de novo* in response to crossovers.

## Discussion

Here, we have shown that phosphorylation of SYP-1 at its PBD-binding motif promotes timely synapsis and progression of meiotic prophase, affects recombination rates, and is critical for early steps in establishing the short and long arm chromosome domains essential for correct meiotic disjunction. SYP-1 phosphorylation begins at the TZ as SYP-1 proteins polymerize between axial elements. Once COs are made and meiocytes enter the late pachytene stage, phosphorylated SYP-1 and PLK-2 localize to short arms and vacate the long arms. This asymmetric distribution is required for the full establishment of functionally distinct short and long arms in diplotene and diakinesis. Prevention of SYP-1 phosphorylation at its PBD-binding motif disrupts PLK-2 relocation from PCs to the SC, delays synapsis and CO formation, and delays the exit from the TZ and early pachytene. Without SYP-1 phosphorylation, meiotic defects from early and late prophase lead to mislocalization of the CPC, resulting in chromosome missegregation and reduced progeny viability. Taken together, our results suggest that PLK-2 acts through SYP-1 (potentially through SYP-1’s PBD-binding motif) to ensure confinement of the chromosome segregation machinery to the short arm subdomain, promoting correct segregation of holocentric chromosomes in meiosis I.

PLK-2 has been shown to bind to PC proteins at nuclear envelope-associated LINC complexes to promote chromosome pairing and synapsis in the transition zone (Harper et al., 2011; Labella et al., 2011). Our results raise the possibility that SYP-1 protein phosphorylated at its PBD-binding motif in the nascent SC could provide a new binding site for PLK-2. This new site could promote meiotic events in two ways: by allowing PLK-2 to move off of PCs (possibly in *cis*), promoting the progression of the cell cycle, as well as by recruiting PLK-2 to the SC to establish short and long arm distinction. PLK-2 bound to SYP-1-phos could then phosphorylate other proteins in the vicinity of the SC, initiating a phosphorylation cascade.

Phosphorylated SYP-1 localizes to the short arm immediately after COSA-1 focus formation in late pachytene, suggesting that this relocation is one of the earliest events to take place after recombination intermediates are committed to COs. Another SC central component, SYP-3, has been shown to dynamically move onto and off of chromosomes during meiotic prophase. In a process dependent on PLK-2, this exchange becomes less dynamic after COs are made (Pattabiraman et al., 2017). Thus, the asymmetric relocalization of phosphorylated SYP-1 coincides with a large-scale reduction in the dynamic properties of the SC, raising the possibility that both of these PLK-2-dependent events are mechanistically connected.

The SYP-1 phosphomutants also affected COSA-1 focus number and crossover recombination rate. In the 12A allele, COSA-1 foci representing CO designation sites departed from the wild-type number of 6 per nucleus, with around 13.5% of nuclei having 5 foci and 14.3% having 7 foci. Perturbation of the SC by partial loss of SYP-1 has been shown to increase COSA-1 number (Hayashi et al., 2010; Libuda et al., 2013); perhaps the alterations in our mutant alleles act in a similar fashion. In contrast, the genetic distance of the *unc-5—dpy-20* interval on chromosome IV increased by more than a factor of two in 12A mutants. The disagreement between the change in COSA-1 foci and the large increase in the genetic map we observed has several non-exclusive possible explanations, including generation of additional crossovers not marked by COSA-1, a shift of crossover formation to a central position (the *unc-5—dpy-20* interval roughly bounds the central third of chromosome IV), or a bias in recovery of chromosomes with crossovers in this region in the surviving progeny.

Loss of SYP-1 phosphorylation nearly always resulted in abnormal persistence of SC or SC-interacting proteins in diplotene and diakinesis, and ICP-1 was never observed at wild type levels in -1 oocytes in *syp-1(12A)* mutants. These very penetrative phenotypes contrast with the relatively high viability (60%) of *syp-1(12A)* mutant self-progeny. If spermatogenesis and oogenesis were equally compromised, this level of viability would suggest that gamete production succeeds at least 77% of the time in the absence of SYP-1 phosphorylation. This would imply that despite the localization defects observed in prophase in *syp-1(12A)* mutants, redundant pathways suffice to deposit CPC components at the short arm by metaphase I in a subset of nuclei. Why some chromosomes in *syp-1(12A)* mutants ultimately succeed in recruiting the CPC to short arms and successfully segregate in meiosis while others do not remains to be determined. One possible explanation is the unpredictable nature of crossover position. In *C. elegans*, the obligatory single CO is most likely to occur in the terminal thirds of chromosomes (Barnes et al., 1995). Since establishment of short and long arm domain patterning appears to be determined by the physical distance from the crossover to the chromosome ends, perhaps the mechanisms behind this functional distinction could be confounded on chromosomes which happen to receive more centrally-located COs. More generally, if COs at different positions (central, arm, or terminal) of chromosomes vary with respect to the ease or speed of subsequent long and short arm domain patterning, then nuclei with an excess of “difficult” COs may be more likely to harbor chromosomes that fail in downstream steps of short and long arm establishment in a sensitized background such as *syp-1(12A)* mutants. Similar reasoning could account for the fact that the *syp-1(10A)* mutant has a slight but significant increase in male production without an increase in embryonic lethality, which suggests the *X* chromosome has a higher nondisjunction rate than autosomes in this mutant. Compared to autosomes, the *X* chromosome has a smaller region of suppressed crossing-over near the chromosome midpoint (Rockman and Kruglyak, 2009), predicting a higher incidence of centrally-located crossovers. Such crossovers, though occurring with low frequency, could result in failure to effectively establish the long and short arm domains in *syp-1(10A)* mutants.

The observation that synapsis is delayed in *syp-1(12A)* or *(T452A)* mutants but not in *syp-1(10A)* mutants suggests that SYP-1 phosphorylation at its PBD-binding domain promotes the timely progression of synapsis. Although immunofluorescence against both native and HA-tagged PLK-2 failed to detect PLK-2 on the SC in the TZ or early pachytene in 10A mutants, a residual amount of PLK-2 at the SC below our detection threshold may suffice to promote timely completion of synapsis in this mutant. However, the fact that many chromosomes achieve timely synapsis even in *plk-2* mutants shows that PLK-2 is not strictly necessary for synapsis. The mechanism of synapsis facilitation by PLK-2 remains unknown, but could involve distributive phosphorylation of SC components, which may alter SC structure or binding affinity. In addition to containing the PBD-binding motif, the C-terminus of SYP-1 interacts with SYP-3 (Schild-Prüfert et al., 2011), raising the further possibility that phosphorylation of SYP-1 at T452 may modulate its binding capacity to SYP-3 and promote assembly of central components.

How could the localization of phosphorylated SYP-1 on the short arm lead to subsequent localization of the CPC on the same domain at the end of meiotic prophase? Our data shows that SYP-1 phosphorylation controls the correct distribution of one of the CPC-guiding histone modifications, phosphorylated histone H3T3. We have shown that SYP-1 phosphorylation is critical for LAB-1 restriction to the long arms, where it normally locally increases the activity of PP1 (Tzur et al., 2012), which we have shown to counteract H3T3 phosphorylation. A previous study in fission yeast demonstrated that the H3T3 kinase Haspin is recruited to centromeres by the cohesin regulator Pds5 (Yamagishi et al., 2010). If this mechanism is conserved in *C. elegans*, we would expect Haspin to be recruited to the axis region of the SC, since cohesins and the *C. elegans* ortholog of Pds5, EVL-14, associate with the lateral elements (Pasierbek et al., 2001; Kim et al., 2014). Although our detection of globally high H3T3ph levels in *gsp-2* mutants shows that PP1 likely suppresses Haspin activity all over chromatin during diakinesis, additional negative reinforcement may be required specifically at the axis region of the long arms where Pds5 may increase the amount of Haspin present. In addition to this potential mechanism of negative regulation on the long arms we describe, previous work has also shown that Haspin is phosphorylated and activated by Polo-like kinase in *Xenopus* extracts (Ghenoiu et al., 2013). PLK-2 constrained to the SC short arm by phosphorylated SYP-1 could similarly activate Haspin, and subsequently enrich the CPC at short arms, in *C. elegans.* Our findings combined with previous results suggest major functions of SYP-1 phosphorylation in guiding the localization of the CPC, by suppressing H3T3 phosphorylation on long arms while promoting it on short arms.

## Acknowledgements

We thank A. F. Dernburg, M. P. Colaiácovo, E. Martinez-Perez, V. Jantsch, A. Villeneuve, A. Woglar, M. Zetka, K. Oegema and A. Severson for antibodies, strains and technical assistance, H. Funabiki and S. Ahmed for critical reading and helpful advice, and the rest of the Carlton lab members for discussion and technical assistance. We also thank the anonymous reviewers whose comments improved this manuscript scientifically and editorially. We are deeply indebted to Y. Kim and A. F. Dernburg for sharing their unpublished results. Many nematode strains were also provided by the Caenorhabditis Genetics Center (CGC), which is funded by the NIH National Center for Research Resources (NCRR). This work was supported by a Japan Society for the Promotion of Science (jsps.go.jp) Postdoctoral fellowship to ASC, and JSPS KAKENHI Grants (No. 24687024 Wakate A, No. 15H04328 Kiban B) to PMC and (No. 15K18477 Wakate B, 17K15064 Wakate B) to ASC. The iCeMS institute was supported by the World Premier International Research Initiative (WPI) of the Ministry of Education, Culture, Sports, Science and Technology, Japan. The authors declare no competing financial interests.

## Author contributions

ASC and PMC conceived the experiments, ASC, CNT, SKC, and TU conducted the experiments, and ASC, CNT and PMC wrote the manuscript.

## Materials and Methods

### *C. elegans* strains and conditions

*C. elegans* strains were grown with standard procedures (Brenner, 1974) at 20°C. Wild-type worms were from the N2 Bristol strain. Mutations, transgenes and balancers used in this study are as follows: *me107[plk-2::HA] (I), plk-2(ok1936)/hT2[bli-4(e937) let-?(q782) qls48](I;III), gsp-2(tm301)/eT1(unc-36)(I;III), meIs8 [Ppie-1::gfp::cosa-1 + unc-119(+)] (II), icmSi44 [Psyp-1::syp-1(T452A) + unc-119(+)](II), icmSi35 [Psyp-1::syp-1(4A) + unc-119(+)](II), icmSi33 [Psyp-1::syp-1(8A) + unc-119(+)](II), icmSi42 [Psyp-1::syp-1(10A) + unc-119(+)](II), icmSi25 [Psyp-1::syp-1(12A) + unc-119(+)](II), icmSi31 [Psyp-1::syp-1(12E) + unc-119(+)] line 2 (II), icmSi32 [Psyp-1::syp-1(12E)+ unc-119(+)] line 3 (II), icmSi28 [Psyp-1::syp-1(12D) + unc-119(+)] (II), icmSi24 [Psyp-1::syp-1(wild type) + unc-119(+)] (II), zim-2 (tm574) (IV), spo-11(me44)/nT1[unc-?(n754) let-? qIs50](IV;V), syp-1(me17)/nT1[unc-?(n754) let-?(m435)] (IV;V), unc-119(ed3) (III);ltIs14[pie-1p::GFP-TEV-STag::air-2 + unc-119(+)], unc-5 (e53), dpy-20(e1282ts) (IV).*

For all mutant analyses, we used homozygous mutant progeny of heterozygous parents.

The *syp-1* phospho-mutants were generated using Mos-SCI (Frøkjær-Jensen et al., 2008) with pCFJ151 plasmids and the strain EG6699. To generate *syp-1 (12A), (12E)* and *(12D)* mutants, the *syp-1* gene fragments were synthesized with respective mutations by Invitrogen GeneArt Strings, and cloned into pCFJ151 using Gibson assembly cloning kit (New England BioLabs). The rest of the *syp-1* phosphomutations were created by either mutating *syp-1 (wt)* or *(12A)* genes cloned in pCFJ151 using PCR-based mutagenesis (for T452A) or by stitching *syp-1 (wt)* and *(12A)* gene fragments by Gibson assembly cloning (for 4A and 8A mutants). At least two transgenic lines were generated for each transgene for mutant phenotype analysis. All *syp-1* transgenes contain the promoter (500bp upstream of the gene) and the 3’ UTR (97 bps downstream of the gene). The insertion of the *syp-1* transgene was verified by DNA sequencing.

### Phosphoproteomics

Wild type N2 and *pph-4.1(tm1598)/hT2[bli-4(e937) let-?(q782) qIs48]* worms were grown on NGM plates containing 25 µg/ml of carbenicillin and 1 mM IPTG (Isopropyl β-D-1-thiogalactopyranoside) spread with HT115 bacteria either carrying an empty RNAi vector (L4440, www.addgene.org/1654) or a *pph-4.1*-RNAi plasmid. To generate the *pph-4.1* RNAi plasmid, the 695 bp region spanning the 2nd, 3rd and 4th exons of *pph-4.1* was amplified from *C. elegans* N2 genomic DNA using primers fw:gctcgtgaaatcctagc and rev:cgaatagataaccggctc flanked by Not1 and Nco1 sites and cloned into L4440. First, N2 and *pph-4.1(tm1598)/hT2[bli-4(e937) let-?(q782) qIs48]* worms (P0) synchronized by starvation were transferred to new plates with food, and worms at the L4 larval stage were harvested in M9 + 0.01% Tween buffer, washed three times with M9 + 0.01% Tween buffer, and distributed to either control or *pph-4.1* RNAi plates. About 30 hours later, these worms on RNAi plates were harvested in M9 + 0.01% Tween buffer and bleached to collect embryos. Collected F1 embryos were distributed to fresh RNAi plates. At time points when these F1 worms were either 1 day post L4 stage or 3 days post L4 stage, half of the F1 plates were exposed to 10 Gy of γ-rays to induce DNA damage. Four hours after irradiation,worms were harvested in M9 Buffer, washed three times with M9 buffer, and frozen at -80C. Two mls of pelleted, frozen worms prepared in this manner were thawed and dissolved in 5 ml of urea lysis buffer (20mM HEPES pH 8.0, 9M urea, 1mM sodium orthovanadate, 2.5mM sodium pyrophosphate, 1mM β-glycerol-phosphate), sonicated for 1 min at 30 sec intervals for 10 times until worm bodies were broken up. The worm lysates were spun down at 20000g for 15 minutes, and supernatants were subjected to PTMScan analysis (Cell Signaling Technology): phosphorylated peptides were enriched by phospho-(Ser/Thr) kinase substrate antibody-immobilized protein A beads, and were analyzed by LC-MS/MS using an LTQ-Orbitrap-Elite, ESI-CID (Thermo Fisher). Phosphoenrichment antibodies used were CST catalog# 9607, 6966, 8139, 8738, 9624, 6967, 5759, 9942, 10001, 9614, 9477, 8134, 2325, 5243, and 3004. Protein assignments were made using Sorcerer. Peptide counts indicated in Supplemental Table 1 show pooled counts from all conditions of worms (± RNAi, irradiation or age) used in this assay.

### Microscopy, Cytology, Antibodies

For all cytological preparations, we followed protocols described in (Phillips et al., 2009). Images were acquired on a Deltavision personalDV microscope (Applied Precision/GE Healthcare) with a CoolSNAP ES2 camera (Photometrics) at 23°C, using 60x PlanApoN 1.42NA or 100x UPlanSApo 1.4NA oil immersion objectives (Olympus), and immersion oil (LaserLiquid, Cargille) at a refractive index of 1.513. The Z spacing was 0.2 µm, and raw images were subjected to constrained iterative deconvolution followed by sub-pixel chromatic shift correction using scripted control of the Priism (Chen et al., 1996) software suite (see code “chromatic-shift” at https://github.com/pmcarlton/deltavisionquant) Image acquisition was performed with the softWoRx suite (Applied Precision/GE Healthcare). Image post-processing for publication was limited to linear intensity scaling and maximum-intensity projection using OMERO (Burel et al., 2015). The following antibodies used in the present study have been described previously: HTP-1(Martinez-Perez et al., 2008), LAB-1(de Carvalho et al., 2008), PLK-2 (Labella et al., 2011), ICP-1(Oegema et al., 2001), HIM-8(Phillips et al., 2005), HTP-3 (MacQueen et al., 2005), SUN-1:Ser8p (Penkner et al., 2009), SYP-1(Harper et al., 2011), ZIM-3 (Phillips and Dernburg, 2006)), phosphorylated SYP-4 (Nadarajan et al., 2017) and COH-3/4 (Severson and Meyer, 2014). Antibodies generated for this work are: rabbit-SYP-1_1Phos antibodies generated using the phosphopeptide [SAPLMTSpTPLTAATR], rabbit-SYP-1_3Phos antibodies generated using the phosphopeptide [SAPLM(pT)S(pT)PL(pT)AATR by Eurofins, all used at 1:100 dilution. All the phosphospecific antibodies were affinity purified using the SulfoLink Immobilization Kit from Thermo Fisher (#44999) using non-phosphorylated and phosphorylated peptides. The following commercial antibodies were used: anti-GFP from Roche (#12600500. 1:500 dilution), anti-Histone H3Thr3ph (Phospho-Histone H3 (Thr3) (D5G1I) mAb) from Cell Signaling Technology (#13576S, 1:10000 dilution), anti-Histone H3Ser10Phos from Active Motif (#39254, 1:1000 dilution), rabbit RAD-51 antibody from SDIX/Novus Biologicals (#29480002, lot# G3048-009A02, 1:1000 dilution), rabbit anti-HIM-3 antibody from SDIX/Novus (#53470002, 1:500 dilution), anti-HA from Covance/BioLegend (#901501, 1:500 dilution), and anti-ZIM-2 from SDIX/Novus (#49270002, 1:500 dilution). Secondary antibodies used were DyLight488, DyLight594, DyLight649, or Alexa-488-conjugated AffiniPure antibodies (1:500 dilution) from Jackson ImmunoResearch. All immunofluorescence was performed on adult worms at 1 day post-L4.

For H3T3 phosphorylation intensity measurements, Fiji (Schindelin et al., 2012) was used to calculate the intensity of H3T3ph on DAPI bodies. Regions of interest (ROI) were drawn based on DAPI-positive pixels in -1 oocyte nuclei, and the average H3Thr3ph pixel intensity within ROIs was measured. To sub-tract background intensity, background regions were drawn in the same nucleus outside of the DAPI bodies, whose average pixel intensity was taken as the background intensity. Intensity data were statistically analyzed by the Mann-Whitney test. The number of DAPI bodies counted for H3T3ph is 151 for the wild type and 111 for *syp-1(12A)* mutants. Quantifications of the length of SUN-1 Ser8ph staining as well as the TZ (defined by clustered nuclei without resolvable chromatids), and of RAD-51 foci were performed, as described in (Sato-Carlton et al., 2014). For quantification of RAD-51 foci per nucleus, the nuclei on the coverslip-proximal side of four gonads were scored for each genotype: the numbers of nuclei scored for zone 1-7 are as follows: for *wt*, 142, 163, 173, 146, 127, 111, 92; for *syp-1*(10A), 210, 241, 172, 129, 142, 104, 104; for *syp-1*(12A), 154, 144, 86, 75, 83, 79, 74.

### Genetic recombination frequencies

Recombination frequencies were calculated as in (Zalevsky et al., 1999): *p*, the map distance, is calculated from the fraction *R* of recombinant self-progeny, as 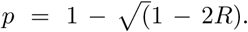 Males of genotype *icmSi25 [Psyp-1::syp-1(12A) + unc-119(+)]*; *syp-1(me17)/nT1[unc-?(n754) let-? qIs50]*, *icmSi44 [Psyp-1::syp-1(T452A) + unc-119(+)]*; *syp-1(me17)/nT1[unc-?(n754) let-? qIs50]* or N2 were crossed with *icmSi25 [Psyp-1::syp-1(12A) + unc-119(+)]; unc-5 (e53), dpy-20(e1282ts); syp-1(me17), icmSi44 [Psyp-1::syp-1(T452A) + unc-119(+)]; unc-5 (e53), dpy-20(e1282ts); syp-1(me17)* or *unc-5 (e53) dpy-20(e1282ts)* hermaphrodites, respectively. Hermaphrodite cross-progeny were picked to single plates, and their progeny were scored for Unc Dpy, wild-type, and Unc non-Dpy or Dpy non-Unc recombinants. The assay was carried out at 23°C to observe the temperature-sensitive phenotype of *dpy-20(e1282ts).* We noted that *syp-1(12A); syp-1(me17)* and *syp-1(T452A); syp-1(me17)* animals generate a small population of sick progeny (worms with abnormal morphology such as arrested development, reduced pigment, early death or Unc): 2.3% and 2.1% of total progeny, respectively, and also *syp-1(12A); syp-1(me17)* animals generate a small number of Dpy progeny: 0.4% of total progeny, presumably due to aneuploidy observed in these mutants. The sick progeny observed in 12A or T452A mutants could be falsely scored as Unc non-Dpy animals in the recombination assay, and indeed we found more Unc non-Dpy animals than Dpy non-Unc in these mutants. Therefore, we limited our analysis to the class of Dpy non-Unc animals to calculate the recombination rate (i.e. multiplied the number of Dpy non-Unc recombinants by two to estimate the true number of recombinants). Since *syp-1(12A); syp-1(me17)* animals were seen to spontaneously generate 0.4% Dpy progeny, we subtracted this percentage from the total number of observed Dpy non-Unc progeny in this cross before calculating.

## Supplemental Materials

**Figure S1.**
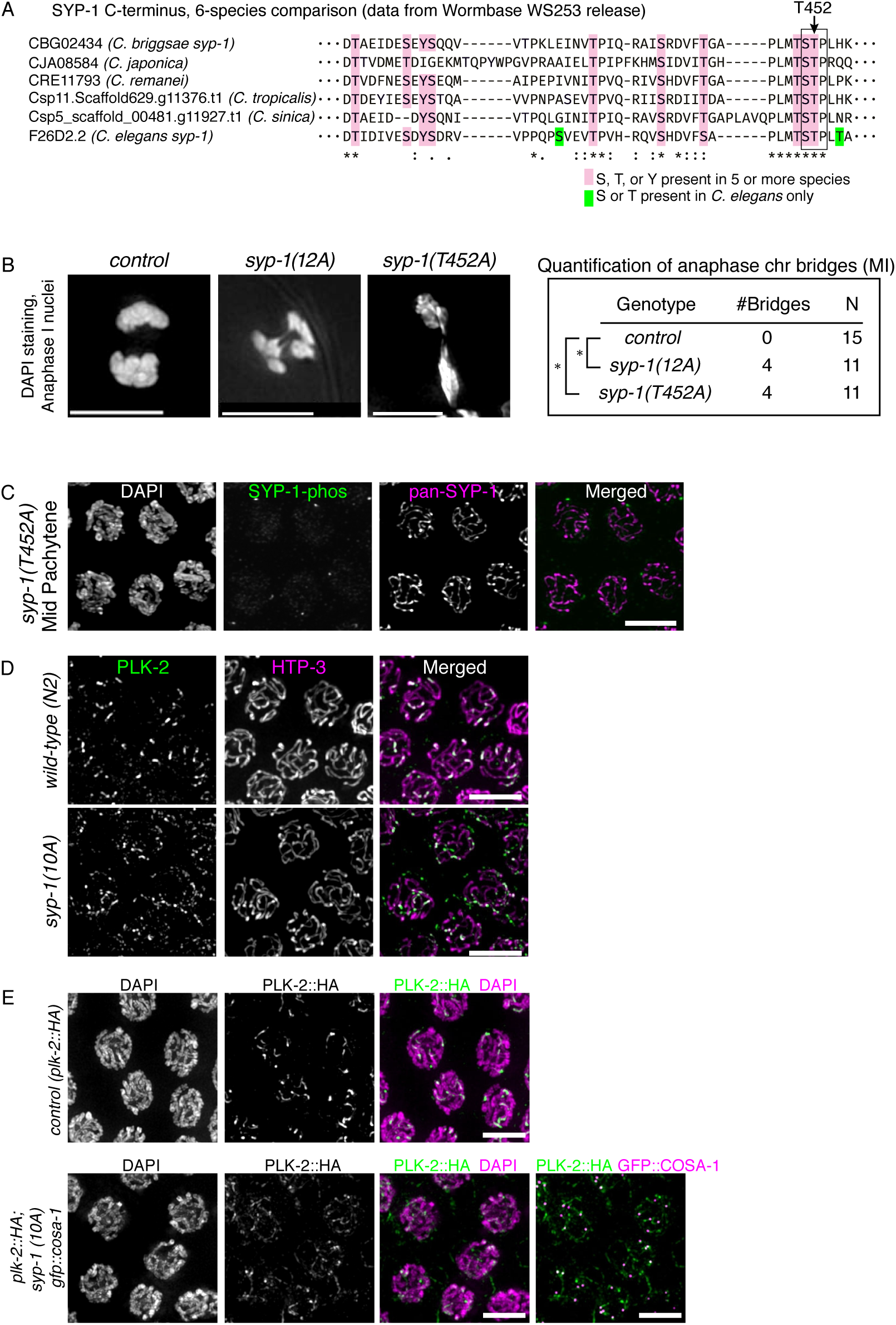
Conservation of the SYP-1 PBD-binding domain among six nematode species, and characterization of *syp-1* phospho mutants. (A) C-terminal amino acid sequence of SYP-1 and its orthologs in six nematode species. Pink highlights Ser, Thr or Tyr residues conserved in 5 or more species; green highlights Ser or Thr only present in *C. elegans*. The conserved PBD-binding motif STP is boxed. (B) DAPI staining images of anaphase I oocyte nuclei in *syp-1(wt); syp-1(me17), syp-1(12A); syp-1(me17)*, and *syp-1(T452A); syp-1(me17)* mutants. Table shows the number of anaphases with visible bridges and the total number of anaphases scored (N). *p=0.02207, Fisher’s exact test. Scale bars, 5 µm. (C) Loss of SYP-1-phos staining in *syp-1(T452A)* mutants. SYP-1-phos is shown in green, and pan-SYP-1 is shown in magenta. Scale bar, 5µm. (D) PLK-2 immunostaining in late pachytene nuclei in wild-type (top) and *syp-1(10A); syp-1*(*me17*) (bottom) gonads. (E) Late pachytene nuclei in control (*plk-2::HA*) and *plk-2::HA; syp-1(10A) gfp::cosa-1; syp-1(me17)* gonads were fixed and stained with antibodies against HA(green), or HA(green) and GFP (magenta) for the 10A mutant. PLK-2::HA is detected at reduced levels on the SC in 10A mutants. Scale bars, 5µm.

**Figure S2.**
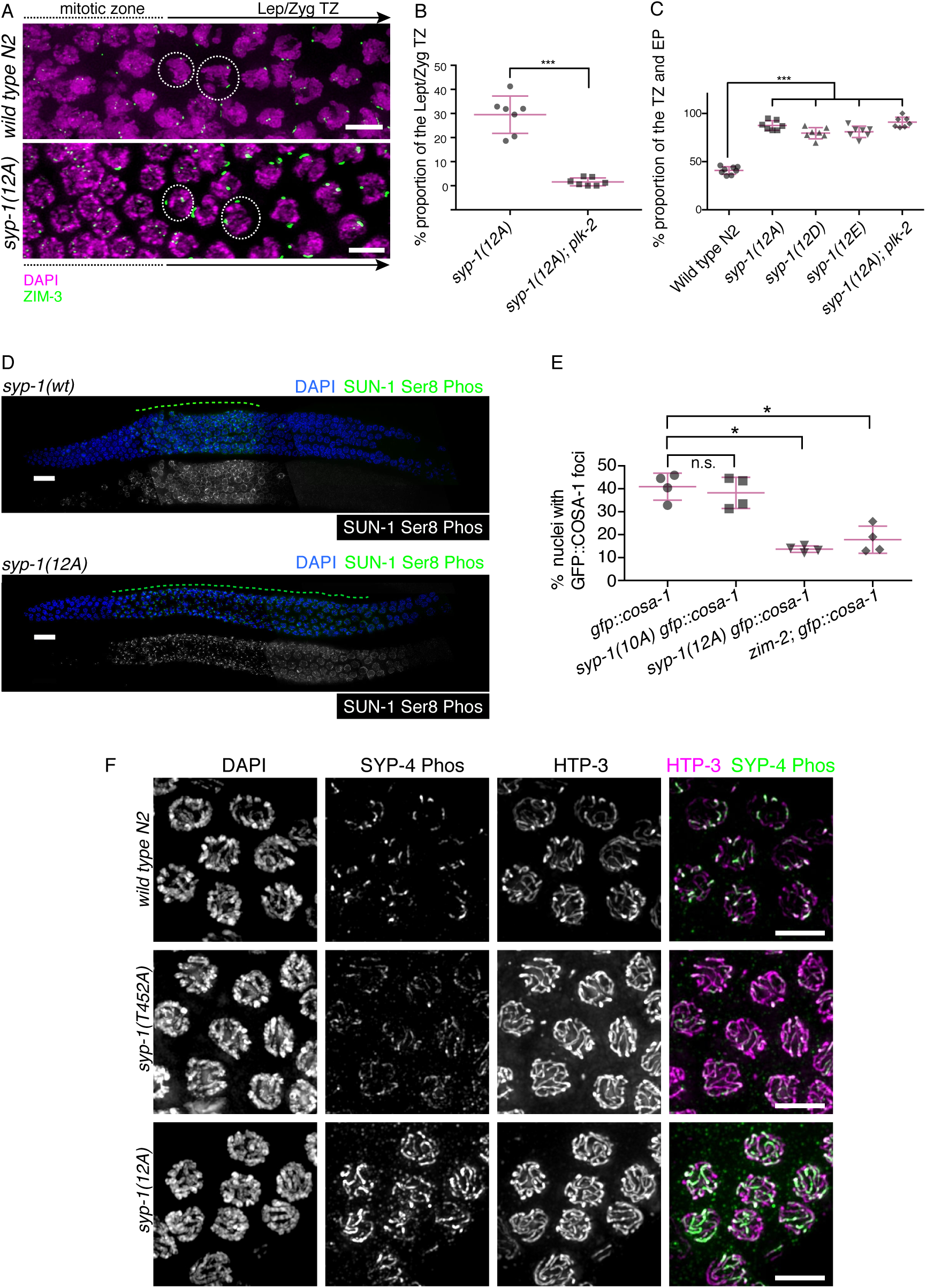
Characterization of *syp-1* phospho-mutants. (A) Homologous chromosome pairing is normal in *syp-1* non-phosphorylatable mutants. Pairing was visualized by immunostaining of ZIM-3 (green), which binds to the PCs of chromosomes I and IV in the wild type and *syp-1(12A); syp-1(me17)* mutant. As meiocytes progress from the zone containing mitotically proliferating cells (left) to meiotic prophase (right), unpaired homologous chromosomes (4 ZIM-3 foci per nucleus) become paired (2 foci per nucleus) in the leptotene/zygotene TZ. DNA is counterstained with DAPI (magenta). Scale bars, 5µm. (B) PLK-2 is required to extend the TZ in *syp-1(12A)* mutants. PLK-2 is essential to induce chromosome clustering, a defining feature of TZ nuclei, and very few or almost no TZ nuclei are found in *plk-2* mutants (Harper et al., 2011; Labella et al., 2011). The proportion of the TZ was calculated by dividing the length of the TZ by the length of TZ + pachytene. The *plk-2(ok1936); syp-1(12A)* double mutant gonads similarly possessed few or no leptotene/zygotene nuclei compared to *syp-1(12A)* controls. ***p<0.001, Mann-Whitney test (C) The proportion of combined TZ and early pachytene region, marked by SUN-1 Ser8 phosphorylation, was extended in *syp-1(12D); syp-1(me17), syp-1(12E); syp-1(me17)* and *plk-2(ok1936); syp-1(12A)* double mutants to a similar level as *syp-1(12A); syp-1(me17)* mutants as well as *plk-2* mutant gonads, previously reported to have an extended region of SUN-1 Ser8 phosphorylation (Harper et al., 2011). ***p<0.0001, Mann-Whitney test. (D) Representative gonad images of *syp-1 (wt); syp-1(me17)* and *syp-1(12A); syp-1(me17)* animals with DAPI (blue) and SUN-1 Ser8-phos staining (green in the top panel and white in the lower panel). The gonad region with SUN-1 Ser8-phos staining is highlighted with a dotted line. Scale bars, 15µm. (E) The appearance of GFP::COSA-1 foci is delayed in *syp-1(12A)* but not in *syp-1(10A)* mutants. Quantification of the proportion of nuclei with GFP::COSA-1 in *gfp::cosa-1, gfp::cosa-1 syp-1(10A); syp-1(me17), gfp::cosa-1 syp-1(12A); syp-1(me17)* and *gfp::cosa-1; zim-2 (tm574)*. The proportion was calculated by dividing the length of the gonad region positive for GFP::COSA-1 by the combined length of the TZ and pachytene. The *zim-2* mutant is defective for homologous pairing of chromosome V, and delays entrance into late pachytene by activating a meiotic checkpoint (Phillips and Dernburg, 2006). This allows *zim-2* to be used as a positive control for delayed GFP::COSA-1 formation. For each genotype, four gonads were scored. Error bars show SD. *p<0.05, Mann-Whitney test. (F) SYP-4-phos immunostaining of late pachytene nuclei in wild type N2, *syp-1(T452A); syp-1(me17),* and *syp-1(12A); syp-1(me17)* mutants. Scale bars, 5µm.

**Figure S3.**
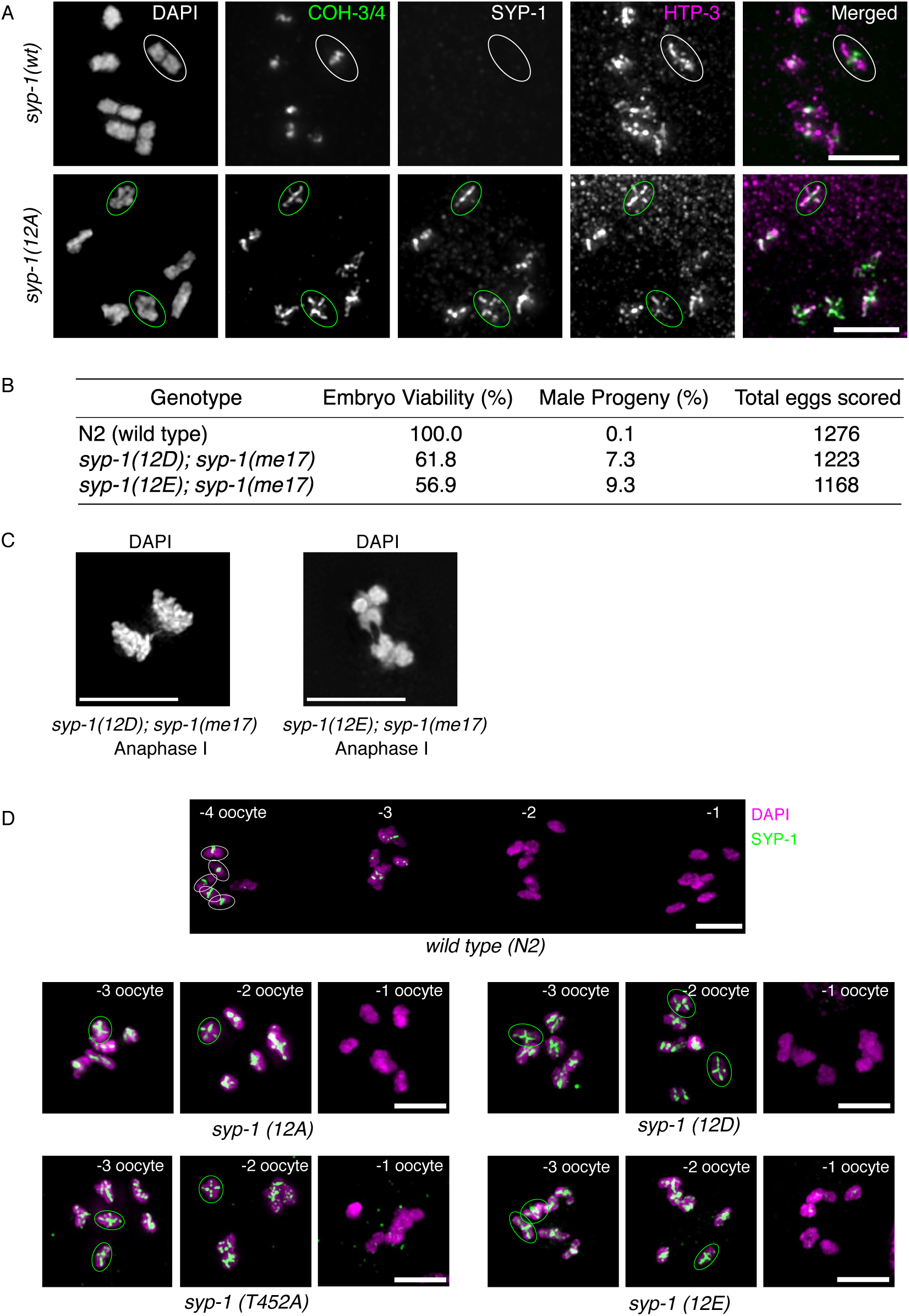
Aberrant persistence of SYP-1 and cohesin subunits COH-3/4 on diakinesis nuclei in *syp-1(12A)* mutants. (A) Immunostaining of -1 oocyte nuclei with COH-3/4 (green), SYP-1 (grayscale), and HTP-3 (magenta) antibodies. COH-3/4 aberrantly persists on both long and short arms in -1 oocyte nuclei in *syp-1(12A); syp-1(me17)* mutants. (B) Percentage of viability and male progeny of worms with the indicated genotypes. For each mutant, all 12 Ser/Thr/Tyr phosphosites are converted to either Asp or Glu. (C) Oocyte anaphase I nucleus with anaphase chromosome bridges in *syp-1(12D); syp-1(me17)* or *syp-1(12E); syp-1(me17)* mutants. Scale bars, 5µm. (D) Pan-SYP-1 staining (green) of diakinesis nuclei in wild-type, *syp-1(12A); syp-1(me17), syp-1(T452A); syp-1(me17), syp-1(12D); syp-1(me17), syp-1(12E); syp-1(me17)* animals. SYP-1 is normally detected on short arms until the -4 or -3 oocyte stage and disappears by the -2 oocyte stage in the wild-type, whereas it always persists on both arms until the -2 or -1 oocyte stage in the *syp-1* phosphomutants indicated. DNA is counterstained with DAPI (magenta). Scale bars, 5µm.

**Figure S4.**
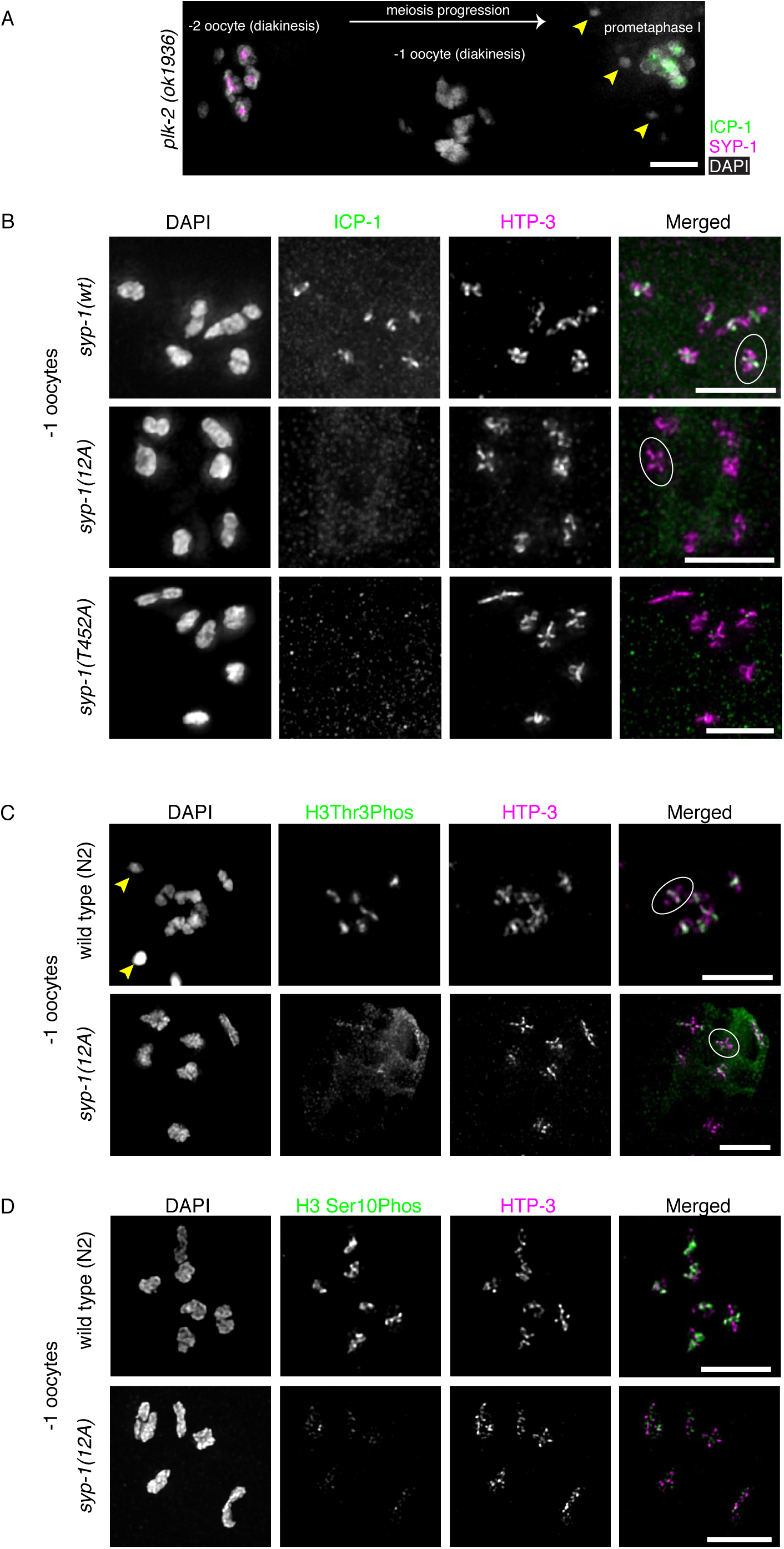
Mislocalization of CPC component or CPC-related histone marks in *syp-1(12A)* mutants. (A) Immunostaining of -2, -1, and prometaphase oocyte nuclei with SYP-1 (magenta) and ICP-1(green) antibodies in the *plk-2 (ok1936)* gonad. Although ICP-1 is not detected in -1 oocyte nuclei, accumulation of ICP-1 is found in oocyte prometaphase nuclei. Arrowheads indicate sperm nuclei. (B and C) Maximum-intensity projection images of the whole -1 oocyte nucleus with ICP-1(green, top) or H3T3ph (green, middle) and HTP-3 (magenta) staining in *syp-1(wt)* (*top*)*, syp-1(me17); syp-1(12A)* (*middle*), and *syp-1(me17)* (*bottom*) (both B and C panels) and *syp-1(T452A); syp-1(me17)* (only panel B) animals. Partial projections of the chromosomes indicated by circles are presented with the long axis rotated to the vertical in Figure 4, panels B and C. Both ICP-1 and H3T3ph staining levels are reduced in 12A mutants. Arrowheads indicate sperm DNA. (D) Immunostaining of histone H3S10ph (green) and HTP-3 (magenta) in -1 oocyte nuclei of wild-type N2 and *syp-1(12A); syp-1(me17)* animals. H3S10ph staining levels are reduced in 12A mutants. Scale bars, 5µm.

**Table S1. Counts of phosphopeptides obtained from mass spectroscopy**

Details of phosphopeptides detected are shown. For all phosphopeptides detected only once, mass spectrometry spectra were manually examined, and peptides with high-confidence assignment are indicated. The MS/MS spectra showing m/z and intensity values for these manually reviewed peptides are shown with accompanying spectrum images in sheets 2 through 8 in the same excel file.

